# Claudin 1-mediated positioning of DC1 to mTECs is essential for antigen transfer-coupled DC1 maturation and maintenance of central tolerance

**DOI:** 10.1101/2025.03.15.643437

**Authors:** Jiří Březina, Tomáš Brabec, David Machač, Matouš Vobořil, Ondřej Ballek, Jan Pačes, Vojtěch Sýkora, Kristína Jančovičová, Evgeny Valter, Katarína Kováčová, Jasper Manning, Valerie Tahtahová, Adéla Čepková, Martina Dobešová, Jan Dobeš, Jan Kubovčiak, Michal Kolář, Petr Kašpárek, Radislav Sedláček, Ondřej Štepánek, Jan Černý, Sachiko Tsukita, Bernard Malissen, Graham Anderson, Dominik Filipp

**Affiliations:** Laboratory of Immunobiology, Institute of Molecular Genetics of the Czech Academy of Sciences, Prague, Czech Republic; Department of Cell Biology, Faculty of Science, Charles University, Prague, Czech Republic; Laboratory of Genomics and Bioinformatics, Institute of Molecular Genetics of the Czech Academy of Sciences, Prague, Czech Republic; Czech Centre of Phenogenomics and Laboratory of Transgenic Models of Diseases, Institute of Molecular Genetics of the Czech Academy of Sciences, Vestec, Czech Republic; Laboratory of Adaptive Immunity, Institute of Molecular Genetics of the Czech Academy of Sciences, Prague, Czech Republic; Advanced Comprehensive Research Organization, Teikyo University, Tokyo, Japan; Centre d’Immunologie de Marseille-Luminy, Aix Marseille Université, Inserm, CNRS, Marseille, France; Institute of Immunology and Immunotherapy, University of Birmingham, Birmingham B15 2TT, UK

## Abstract

The mechanisms of central tolerance, which rely on the presentation of self-antigens by medullary thymic epithelial cells (mTECs) and DCs, prevent autoimmunity by eliminating self-reactive T-cells. While mTECs produce self-antigens in an autonomous manner, DCs acquire them from mTECs via cooperative antigen transfer (CAT). Our recent data showed that preferential pairing occurs between distinct subsets of mTECs and DCs in CAT, providing a rationale for the existence of molecular determinants which control such pairing and the outcome of central tolerance. Here, we compared the transcriptomes of CAT-experienced and -inexperienced DCs and identified Claudin 1 as a molecule involved in CAT-coupled type 1 DC (DC1) maturation. By mapping thymic DC1 heterogeneity, we identified their early and late maturation states. DC1-specific ablation of Claudin 1 led to a reduction in CAT-experienced late mature DC1s and hampered DC1 maturation. These phenotypes correlated with the displacement of DC1s from the vicinity of mTECs. This translated into impaired Treg selection and clonal deletion of TRA-specific T-cells manifested via a break in tolerance and symptoms of multi-organ autoimmunity. Collectively, our results identify thymic DC1-derived Claudin 1 as a regulator of immune tolerance.

**One Sentence Summary:** The expression of Claudin 1 on type 1 dendritic cells regulates their proximity to mTECs, which is required for effective antigen transfer coupled with DC1 maturation and establishment of T-cell tolerance.

## INTRODUCTION

The vast T-cell receptor (TCR) repertoire of the adaptive immune system would be detrimental to the host if self-reactive T-cells were not properly eliminated (*1*). The mechanistic basis for this selection, which occurs largely in the thymic medulla, is the presentation of self-antigens to developing T-cells, a process known as central tolerance. Medullary thymic epithelial cells (mTECs) and thymic dendritic cells (DCs) are antigen presenting cells (APCs) which are instrumental in this process (*2*, *3*). In addition to self-antigens which are expressed ubiquitously, the murine genome encodes approximately 6500 genes whose products, referred to as tissue-restricted antigens (TRAs) (*4*), e.g. insulin (*5*) or enteric defensins (*6*), are only found in a limited number of extrathymic tissues (*1*). To prevent TRA-targeted autoimmunity, TRAs are also expressed on mTECs, which employ an unique transcriptional machinery that is directed by a unconventional transcriptional modulator, Autoimmune regulator (Aire), that mediates ectopic TRA expression (*7*). Fragments of TRAs, as well as those derived from other generic antigens, are directly presented on mTEC MHC molecules that are recognized by developing self-reactive T-cells, leading to clonal deletion (recessive tolerance) or conversion to T regulatory cells (Tregs) (dominant tolerance) (*2*). Interestingly, thymic DCs do not express Aire but instead present TRAs indirectly through TRA acquisition from mTECs (*8–10*). This process of directional antigen spreading is referred to as cooperative antigen transfer (CAT) and has been shown to be essential in the reinforcement of both recessive and dominant tolerance (*3*, *11*, *12*).

A subset of mTECs that expresses high levels of MHCII, CD80/86, AIRE, and TRAs is referred to as mTEC^HI^, while another subset, mTEC^LO^, exhibits low expression of the same molecules and limited TRA expression (*13*). Regarding mTEC^HI^ it has been recently shown that these cells give rise to a variety of cell types that mimic the transcriptome and phenotype of differentiated peripheral cells such as keratinocytes, tuft cells, and microfold cells. These “mimetic cells” serve alongside mTEC^HI^ as a central reservoir of TRAs for CAT (*12*, *14*, *15*).

Thymic DCs with their potential to acquire mTEC-derived antigens, are represented by two conventional DC lineages, type 1 (DC1) and 2 (DC2) (*16–18*). The cells of both lineages are efficient in the acquisition of mTEC-derived antigens but differ significantly in their mTEC subset preferences. In particular, DC1s prefer to uptake antigens from mTEC^HI^ and mimetic cells which are loaded with TRAs, while the DC2 lineage interacts preferentially with mTEC^LO^ (*10*). It is of interest that monocyte-derived thymic CD11c^+^ cells complement DC1 and DC2 by being effective in CAT not only from various mTEC subsets but also from other CD11c^+^ cells (*10*, *19*).

Recent studies have described CAT as a complex and highly organized process in which preferential engagement of specific mTEC and DC subsets suggests a deterministic nature of their interaction (*9*, *10*, *16*, *20–25*). However, the molecules which drive mTEC-to-DC CAT are largely unknown, with the exception of the scavenger receptor CD36, that mediates the terminal phase of CAT, i.e. the “scavenging” of mTEC-derived apoptotic bodies by DC1s (*22*). In this context, CD36 is required for the transfer of surface but not cytoplasmic molecules by CAT, which is indicative of trogocytosis. However, we recently found that the transfer of membrane-bound proteins is a less frequently observed mechanism of CAT to DC1s in comparison to the transfer of cytoplasmic molecules, i.e. the phagocytosis of mTEC apopototic bodies (*10*).

Recently, it has been also shown that the engulfment of apoptotic cells within tumors and peripheral organs drives homeostatic maturation of DC1s to an activated cell state known as aDC1 (*26–29*). aDC1s possess superior antigen presentation that is accompanied by transcriptomic changes in cholesterol metabolism (*27*, *30*). Interestingly, several studies have provided evidence that immature DC1s give rise to aDC1s in the thymus (*18*, *30–32*). Hence, our intention was to determine the molecule(s) which are involved in CAT and/or CAT-coupled DC1 maturation in the thymus.

In this study, we identified the tight junction protein CLAUDIN 1, encoded by the *Cldn1* gene, which impacts antigen transfer-coupled DC1 maturation. Comparative analysis of single-cell RNA sequencing (scRNAseq) of CAT-experienced and -inexperienced myeloid thymic cells allowed us to design a flow cytometry-based gating strategy to redefine the heterogeneity of thymic DCs and their distinct maturation states. Analysis of 3D-light sheet fluorescence microscopy images of the thymic medulla determined that Claudin 1 positions DC1s in direct contact with mTECs. As part of this study, we utilized a *Defa6^iCre^R26^TdTOMATO^*mouse model in which TdTOMATO was expressed in mTECs through Aire-dependent activation of the *Defa6* promoter, thus mimicking the expression of the natural TRA, enteric α-defensin 6 (*6*, *10*). Infusing bone marrow (BM) from a mouse harboring a conditional deletion of Claudin 1 in the DC1 lineage (*33*, *34*) into this model, we determined that Claudin 1 is critical for the presence of CAT-experienced late mature DC1s, ensuring proficient indirect TRA presentation to self-reactive T-cells. Indeed, by constructing the novel mouse model, *Defa6^iCre^R26^TdT-OVA^*, we demonstrated the impact of Claudin 1 on Treg selection and clonal deletion of TRA-specific T-cells which subsequently resulted in a break in tolerance.

## RESULTS

### Claudin 1 is upregulated in CAT-experienced DC1s

To reveal molecules involved in CAT, we compared scRNAseq data of CAT-experienced and CAT-inexperienced myeloid cells from the thymus of 6 week old *Foxn1^Cre^R26^TdTOMATO^* mice, in which the production of TdTOMATO was restricted to TECs (Fig. 1A, S1A) (*10*, *19*). Since thymic myeloid cells exhibit significant heterogeneity, we first determined their composition (Fig. S1B). We excluded from this analysis the following cell populations: granulocytes (Gran; *Ly6g*), T, B, and NK cells (T B NK; *Lck*, *Cd79a*, *Klrb1c*), T-APC doublets (*Lck*, *H2-Aa*) and pDCs (*Siglech*), all of which are not (or marginally) involved in CAT (Fig. S1B-C) (*10*, *21*). Three distinct lineages of myeloid APCs were found: (i) DC1, (ii) DC2, and (iii) monocyte/macrophage (Mono, Mac) (Fig. 1B). We observed previously undescribed heterogeneity regarding their maturation states. Among DC1 and DC2 lineages, we detected *Ccr7^−^* immature DCs (DC1/2), some of which were proliferating *Mki67^+^* cells (DC1/2 prolif) and *Ccr7^+^* mature DCs (aDC1/2), the latter which have been described as being in early or late mature state (*18*, *27*, *30*, *35*). In this study, we refer to early and late as the “a” and “b” maturation states, respectively (Fig. 1B). Accordingly, while in the “a” state, the lineage specific markers of DC1 and DC2, *Xcr1* and *Sirpa,* respectively (Fig. S1D-F) were readily detectable, the “b” state of these lineages exhibited a profound decrease in the expression of these markers. Surprisingly, in our scRNAseq analysis, we found an additional DC cluster which transcriptionally corresponded to the “b” state, however due to the lack of both *Xcr1* and *Sirpa* markers, its origin in respect to DC1 or DC2 lineages was unclear. Since this *Xcr1*/*Sirpa* double negative population clustered with aDC1a (Fig. 1B), we designated this cluster as aDC1b. In addition, our scRNAseq analysis revealed high expression levels of *Cd81* and *Il7r* in aDC1b and aDC2b, respectively, which we consequently used as specific markers for these subsets (Fig.S1E-F).

**Figure 1.**
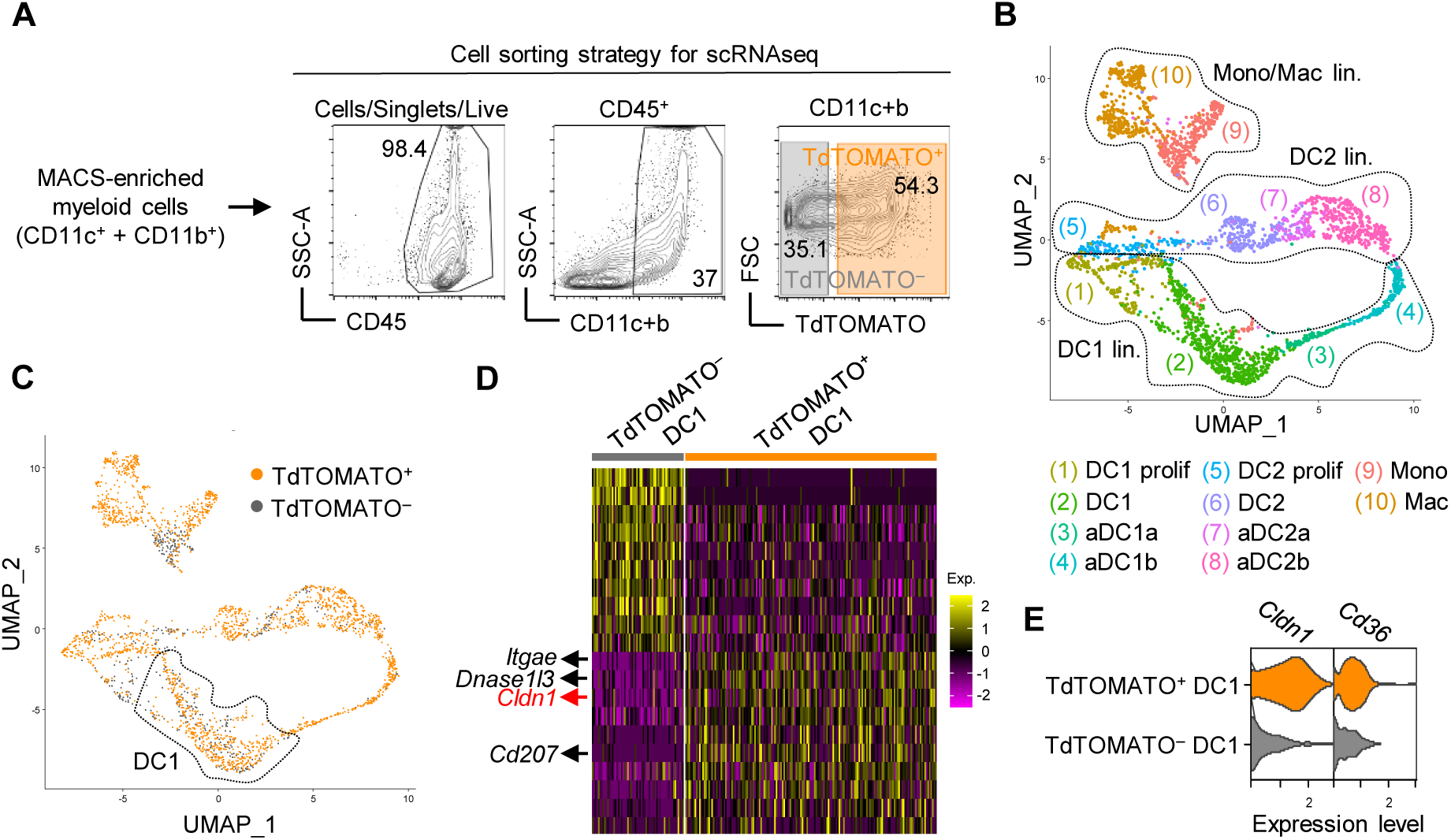
Claudin 1 is upregulated in CAT-experienced DC1s. (**A)** Cell sorting gating stategy used to perform scRNAseq of thymic myeloid cells. Thymic cells were isolated from *Foxn1^Cre^R26^TdTOMATO^* mice, MACS-enriched for CD11c^+^ and CD11b^+^ cells, and sorted as either as TdTOMATO^+^ or TdTOMATO^−^ CD45^+^ CD11c^+^/CD11b^+^ cells. **(B)** UMAP of annotated thymic myeloid cells from scRNAseq excluding subsets shown in red in the legend of Fig. S1B. Individual cell lineages are demarcated by dotted lines. DC=conventional DC, aDC=activated DC, Mac=Macrophages, Mono=Monocytes, prolif=proliferating. **(C)** UMAP of scRNAseq corresponding to Fig. 1B projecting CAT-experienced (orange) and CAT-inexperienced (grey) cells. Dotted line shows the DC1 subset. **(D)** Heatmap showing top 10 down- and up-regulated genes in CAT-experienced over CAT-inexperienced DC1s. Heatmap color scale depicts average log2 fold change. **(E)** Violin plots show the expression of *Cldn1* and *Cd36* by CAT-experienced (orange) and CAT-inexperienced (grey) DC1.

Using selected markers that were identified by scRNAseq (Fig. S1F), we designed a flow cytometry gating strategy that separated both DC lineages from the monocyte/macrophage lineage (Fig. S2A) based on the combinatorial expression of *Cd14, Sirpa, and Mgl2* (Fig. S1F). As expected, CCR7^+^ aDCs were comprised of aDC1a, aDC1b, aDC2a, and aDC2b, while CCR7^−^ DCs consisted of immature DC1 and DC2 subsets (Fig. S2A-D). In a seminal study, a set of genes referred to as “MAT ON genes” was found to be associated with homeostatic maturation of the thymic DC1 lineage (*30*). Remarkably, these genes which encode for chemokines, costimulatory, and checkpoint molecules, i.e. *Ccl17, Ccl22, Cd40* or *Cd274* are critical in mediating clonal deletion and Treg selection (*31*, *36*, *37*). Our scRNAseq analysis confirmed that MAT ON genes are upregulated during maturation of DC1 to aDC1a (Fig. S2E) (*30*). However, we detected a more pronounced upregulation of these genes in the aDC1b subset which fits with the existence of a late mature state of the DC1 lineage (Fig. S2E). Consistent with this notion is the fact that both aDC1a and aDC1b expressed high levels of genes involved in MHCII antigen presentation such as *H2-Eb2*, which was expressed exclusively by aDC subsets (Fig S2). It is of note that we also observed the same expression profile of both MAT ON genes and genes involved in MHCII antigen presentation in the DC2 lineage (Fig. S2E-F).

We recently established that the DC1 lineage is the most effective in CAT (*10*). In addition, it has been shown that the thymic DC1 but not the DC2 lineage is specialized for TRA acquisition (*10*, *16*, *22*), which is vital for clonal deletion and Treg selection to occur (*2*). Thus, we set out to identify molecules that affect CAT and/or processes triggered by CAT. Since we hypothesized that gene products that are involved in CAT would be upregulated only in CAT-experienced cells, we conducted our scRNAseq on both CAT-experienced (TdTOMATO^+^; orange) and CAT-inexperienced (TdTOMATO^−^; grey) cells (Fig 1C) and identified the most up-regulated genes in TdTOMATO^+^ versus TdTOMATO^−^ cells from the DC1 lineage (Fig. 1D). Since we observed that CAT occurs in immature DC1s (*10*), we directed our search for determinants within this subset (Fig. 1D). Among the top upregulated genes in CAT-experienced DC1s were *Itgae* (CD103), *Cd207* (Langerin), and *Dnase1l3* (Fig. 1D) all of which have been found to be involved in the engulfment of apoptotic cells (*27*, *38*, *39*). Among this group of genes, we also identified *Cldn1* (Fig. 1D), which encodes the tight junction protein CLAUDIN 1 (*40*). Intriguingly, Claudin 1 has been implicated in the process by which DC1s interact with the intestinal epithelial layer to acquire microbial antigens in the context of the immune response (*41*, *42*), a mechanism that has been also described in other epithelial organs such as the lungs and skin (*43*, *44*). Because mTECs form an epithelial network that is connected by tight junctions (*45*, *46*), we predicted that in the context of immune tolerance, Claudin 1 mediates the interaction of DC1s with mTECs. In fact, while the expression of the scavenger receptor *Cd36* showed only mild enrichment in TdTOMATO^+^ DC1, the enrichment of *Cldn1* was far more dramatic (Fig. 1E).

### Lineage tracing and CLAUDIN 1 protein expression in thymic DC1s

Since our scRNAseq analysis revealed the presence of CCR7^+^XCR1^−^SIRPA^−^ DCs, (denoted as aDC1b) whose ontogeny was unclear (Fig. S1D-F, S2A, S2C-D), we fate-mapped thymic DCs using *XCR1^iCre^R26^TdTOMATO^* mice (*34*, *47*), in which TdTOMATO^+^ cells represented DCs with a history of XCR1 expression (Fig. 2A). Indeed, all thymic DC1 and aDC1a were TdTOMATO^+^ in *XCR1^iCre^R26^TdTOMATO^*mice (Fig. 2B, S3A). Remarkably, approximately 90% of aDC1b were also TdTOMATO^+^ demonstrating that aDC1b which have lost XCR1 expression are members of only the DC1 lineage.

**Figure 2.**
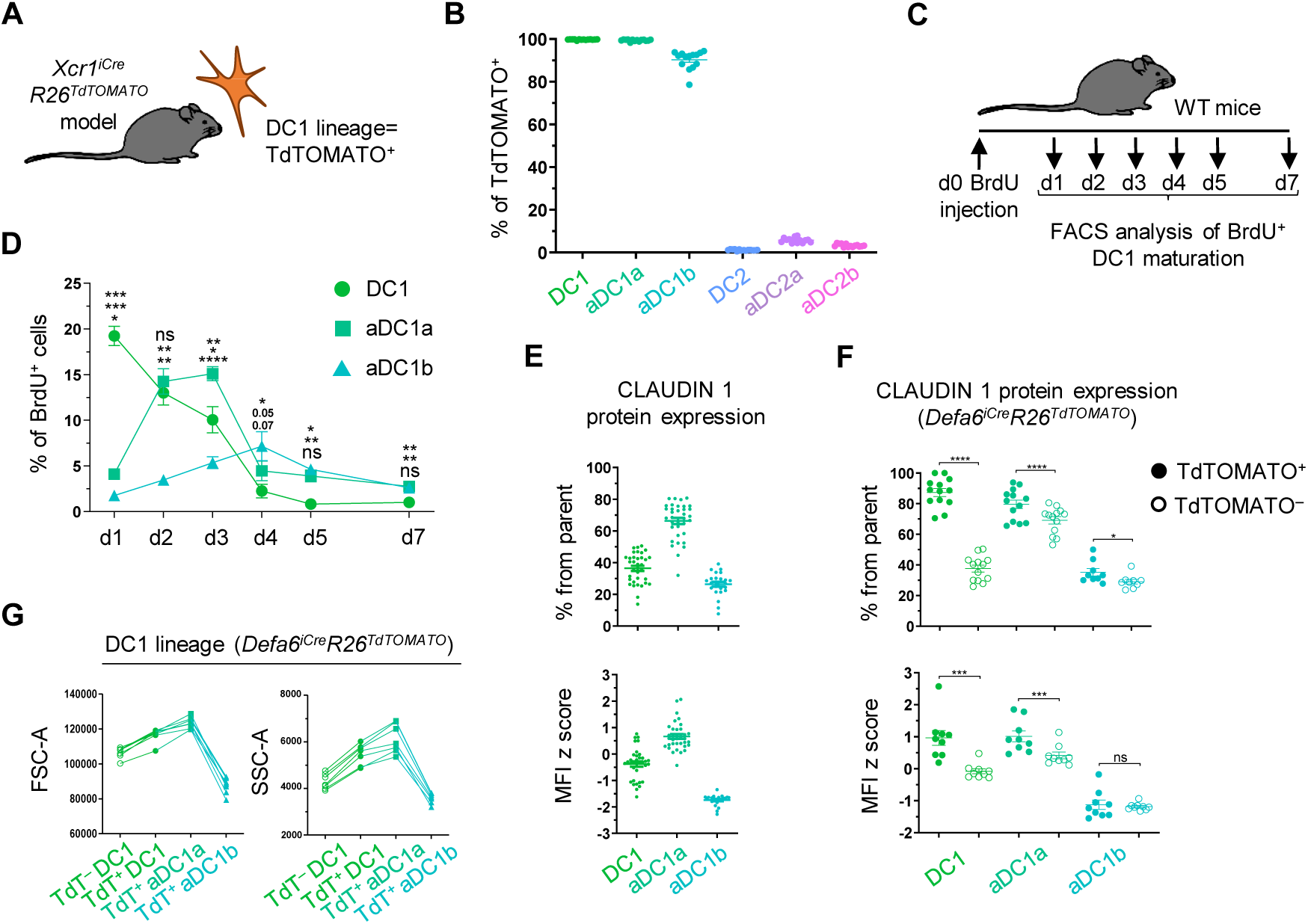
Lineage tracing and CLAUDIN 1 protein expression in thymic DC1s. (**A**) Schematic of mouse model used for lineage tracing. (**B**) Frequency of TdTOMATO^+^ cells within DC subsets from Fig. S3A (mean ± SEM, *n*=14 mice from 3 independent experiments). (**C**) Design of BrdU DC1 lineage tracing experiment. (**D**) Percent of BrdU^+^ cells within DC1 lineage subsets on indicated days after BrdU administration related to Fig. S3B (mean ± SEM, *n*=3-5 mice from 2 independent experiments). (**E**) The frequency of CLAUDIN 1^+^ cells and MFI z score of CLAUDIN 1 expression within DC1 subsets (mean ± SEM, *n*=24-35 mice from a minimum of 4 independent experiments). **(F)** Frequency of CLAUDIN 1^+^ cells and MFI z score of CLAUDIN 1 expression within TdTOMATO^+^ and TdTOMATO^−^ DC1 subsets from thymi of *Defa6^iCre^R26^TdTOMATO^*mice (mean ± SEM, n=9-14 mice from a minimum of 3 independent experiments). **(G)** Graphs showing forward scatter area (left panel) and side scatter area (right panel) of TdTOMATO^−^ (TdT^−^) and TdTOMATO^+^ (TdT^+^) subsets from DC1 lineage isolated from thymi *Defa6^iCre^R26^TdTOMATO^* mice (mean ± SEM, *n*=7 mice from 3 independent experiments). Statistical analysis in D was performed using RM one-way ANOVA with Tukeýs multiple comparisons test and in E using paired, two-tailed Student’s t-test, p ≤ 0.05 = *, p ≤ 0.01 = **, p ≤ 0.001***, p < 0.0001 = ****, ns = not significant.

To determine if aDC1b cells are more mature than DC1 or aDC1a, we injected WT mice with BrdU and analyzed its redistribution in defined subsets of the DC1 lineage over a seven-day period (Fig. 2C). Given that only immature DC1s are capable of proliferation (Fig. 1B, S1F) (*35*), only this subset incorporated BrdU after one day. In the ensuing days, the frequency of BrdU^+^ DC1s gradually decreased (Fig. 2D, S3B) while the frequency of BrdU^+^ aDC1a and aDC1b peaked at day three and day four, respectively. This is consistent with a scenario whereby DC1 gives rise to early mature aDC1a, a portion of which continues to develop towards late mature aDC1b. When CLAUDIN 1 protein expression was assessed across these subsets, aDC1a exhibited the highest level while aDC1b the lowest (Fig. 2E, S3C).

Several studies have reported that CLAUDIN 1 forms tight junctions via heterotypic binding to CLAUDIN 3, which has been shown to be a lineage marker of Aire-expressing mTECs (*48*, *49*), Therefore, we hypothesized that Claudin 1 binding to Claudin 3 would juxtapose DC1s to mTECs. As a starting point, we first assessed the levels of Claudin 3 expression across various thymic and mTEC subsets including mTEC^LO^, mTEC^HI^, Ly6D^+^ (keratinocyte mimetics), and Ly6D^−^ mimetic cells (Fig. S4A). We also analyzed the mTEC subset that expresses a prototypical Aire-dependent TRA, α-Defensin 6 (Defa6^+^ mTECs) in our *Defa6^iCre^R26^TdTOMATO^ model* (*10*). Our results were consistent with the findings of *Hamazaki et.al.* (*50*), that mTECs had the highest frequency of Claudin 3 expression (Fig. S4B-C) with Aire-independent mTEC^LO^ having the lowest frequency of all mTEC subsets analyzed (Fig. S4D-E). Importantly, increasing levels of Claudin 3 expression were detected along the developmental pathway of Aire^+^ mTEC^HI^ into Aire^−^ mimetic cells suggesting that TRA-loaded mTEC subsets were the main target for CAT by the Claudin1^+^ DC1 subsets. In line with this observation, we detected the highest expression of Claudin 3 in Defa6^+^ mTECs and Ly6D^+^ mimetic cells (Fig. S4D-E).

Since DC1, aDC1a, and aDC1b are members of the thymic DC1 lineage, we compared their ability to acquire TRAs in the *Defa6^iCre^R26^TdTOMATO^*mouse model. The highest frequency of TdTOMATO was detected in aDC1a, an intermediate frequency in aDC1b, and the lowest frequency in DC1 (Fig. S5A-B). In agreement with our scRNAseq, TdTOMATO^+^ DC1s showed a higher frequency of CLAUDIN 1^+^ cells as well as a higher level of CLAUDIN 1 expression in comparison to TdTOMATO^−^ cells (Fig. 2F). This indicated that the initiation of CAT to DC1s triggered strong upregulation of CLAUDIN 1 expression. In fact, nearly all TdTOMATO^+^ DC1 and aDC1a analyzed expressed CLAUDIN 1. Importantly, as depicted in Fig. 2G, the initiation of CAT correlated with the continuous increase in the size and granularity of DC1s until they reached aDC1a, at which point the values associated with these parameters were dramatically decreased below what was observed in TdTOMATO^−^ DC1s. Therefore, we concluded that an intensive engulfment of apoptotic bodies by CAT occured chiefly between DC1 and aDC1a. Once the cells are satiated, CAT precipitously wanes while the ensuing maturation to aDC1b is marked by antigen processing and presentation (Fig. S2E-F).

### Claudin 1 is involved in CAT-coupled homeostatic DC1 maturation

In order to determine if Claudin 1 is a molecular determinant of CAT and/or CAT-coupled maturation of the DC1 lineage, we ablated its expression by crossing a *Cldn1^fl/fl^* mouse strain (*33*) with a *XCR1^iCre^* mouse (*34*). In *XCR1^iCre^Cldn1^fl/fl^* mice, Claudin 1 was not detected in the DC1 lineage (Fig. S5C-D) and the frequency of individual DC1 subsets remained unchanged in comparison to controls (Fig. S5E). To analyze CAT of TdTOMATO protein, *Defa6^iCre^R26^TdTOMATO^*mice were sub-lethally irradiated and reconstituted with a mixture of BM from *XCR1^iCre^Cldn1^fl/fl^* and WT mice at a 1:1 ratio (Fig. 3A, S5F-G). This experimental setup enabled us to compare the participation of Claudin 1-sufficient versus -deficient DC1 lineage in CAT within a shared thymic microenvironment. We observed that the median fluorescence intensity (MFI) of TdTOMATO was comparable between Claudin 1-sufficient and -deficient cells in all three DC1 lineage subsets (Fig. S5H) suggesting that Claudin 1 did not affect the intracellular processing of acquired antigens.

**Figure 3.**
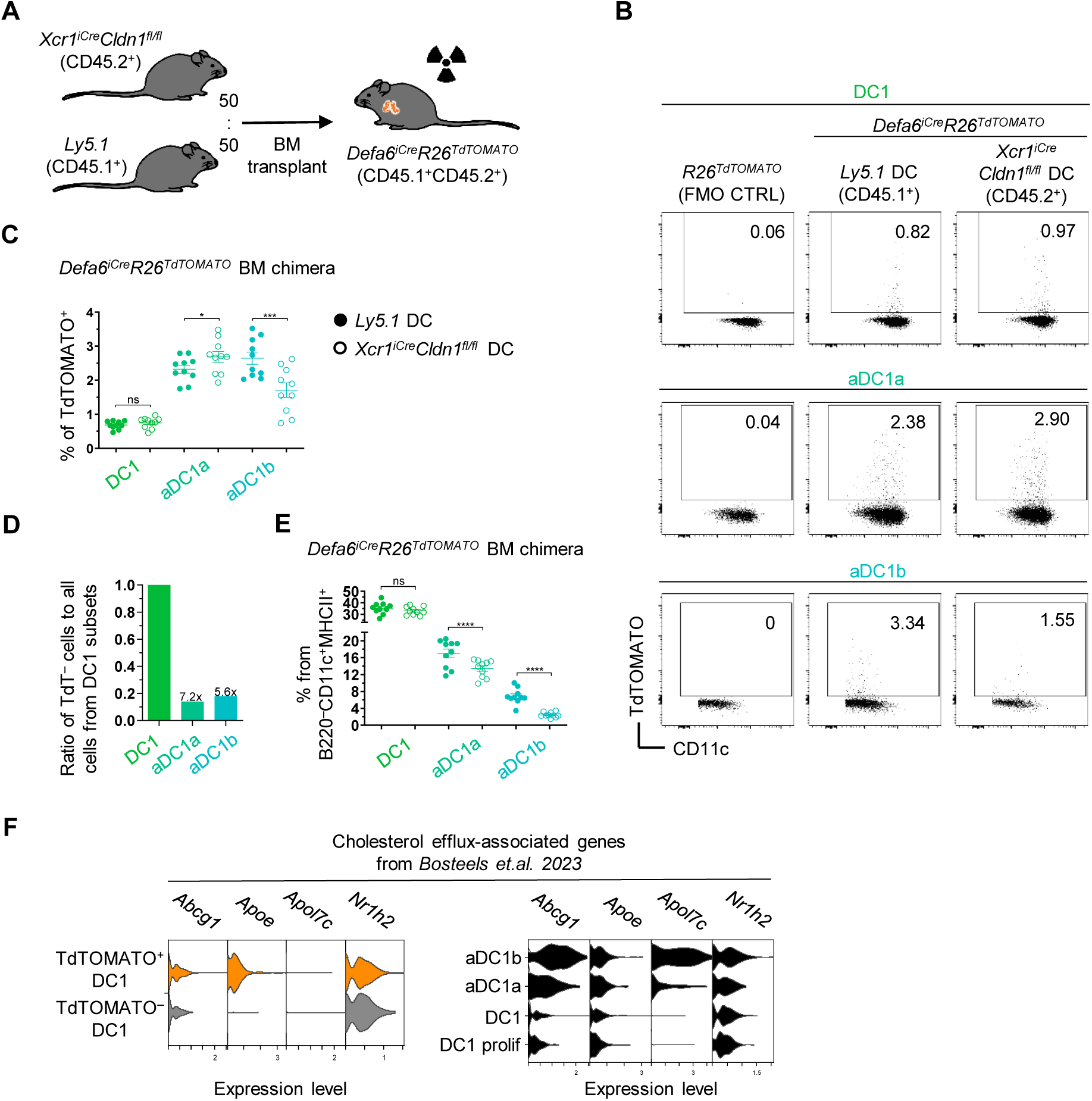
Claudin 1 is involved in CAT-coupled homeostatic DC1 maturation. **(A)** Schematic of competitive BM chimera experiment assessing the role of Claudin 1 in CAT. **(B)** Representative flow cytometry plots show the frequency of TdTOMATO^+^ cells within Claudin 1-sufficient (*Ly5.1* BM) and -deficient (*XCR1^iCre^Cldn1^fl/fl^* BM) DC1 subsets from from competitive BM chimeras in Fig. 3A. FMO controls are shown (*R26^TdTOMATO^* mouse). **(C)** Frequency of TdTOMATO^+^ cells within Claudin 1-sufficient and -deficient DC1 subsets from Fig. 3B (mean ± SEM, *n*=10 mice from 3 independent experiments). **(D)** Ratio of TdTOMATO^−^ (TdT^−^) cells to all cells within each indicated DC1 lineage subset (i.e. DC1, aDC1a and aDC1b) taken from scRNAseq (Fig. 1). These ratios were normalized to the ratio in DC1 subset which was given the reference value “1“. The number above the aDC1a and aDC1b bars indicates the fold change reduction in the ratio value in comparison to DC1 subset. **(E)** Frequency of individual DC1 subsets within thymic DCs from Claudin 1-sufficient (solid circle) and -deficient (empty circle) BM from Fig. 3A (mean ± SEM, *n*=10 mice from 3 independent experiments). **(F)** Violin plots from scRNAseq analysis (Fig. 1) show the expression of cholesterol efflux-associated genes within CAT-experienced (orange) and CAT-inexperienced (grey) DC1 (left panel) and DC1 lineage subsets (right panel). Statistical analysis in C, and E was performed using paired, two-tailed Student’s t-test, p ≤ 0.05 = *, p ≤ 0.001***, p < 0.0001 = ****, ns = not significant.

When compared to WT TdTOMATO^+^ cells, we found a pronounced decrease in the frequency of Claudin 1-deficient TdTOMATO^+^ aDC1b. This result was unexpected since this decrease was not observed in Claudin 1-deficient TdTOMATO^+^ DC1s and aDC1a (Fig. 3B-C), which are the main executors of CAT (Fig. 2G). Since aDC1b participation in CAT is limited (Fig. 2G), we redirected our attention to the possibility that CLAUDIN 1 is involved in CAT-coupled maturation. In this context, previous studies demonstrated that DC1 maturation is coupled to the engulfment of apoptotic bodies (*26–29*). Our data from scRNAseq is in line with this conclusion (Fig. 1C). Specifically, the ratio of CAT-inexperienced (TdTOMATO^−^) to CAT-experienced (TdTOMATO^+^) aDC1a and aDC1b was 7.2 and 5.6 times lower, respectively, in comparison to the same ratio in the DC1 subset (Fig. 3D). Since CAT-inexperienced cells exhibited a limited capability to undergo maturation, CAT appears to be a prerequisite for DC1 maturation in the thymus. Furthermore, the small number of TdTOMATO^−^ cells found in the aDC1a and aDC1b subsets (Fig. 1C) may be due to TdTOMATO degradation. Next, we assessed if the absence of Claudin 1 would affect the maturation of thymic DC1s in *Defa6^iCre^R26^TdTOMATO^* BM chimeric mice (Fig. 3E). Consistent with the involvement of CLAUDIN 1 in CAT-coupled DC1 maturation, while there was no effect of Claudin 1 deficiency on the percentage of immature DC1s, the overall frequency of aDC1a was significantly reduced with an even higher reduction (nearly 3-fold) in the percentage of aDC1b in comparison to controls.

Regarding a mechanism that is responsible for homeostatic maturation, it has been shown that cholesterol is a principal component of apoptotic bodies which drives the maturation of splenic DC1s. This process is accompanied by the upregulation of cholesterol efflux-associated genes, such as *Apoe, Abcg1, Apol7c and Nr1h2* (*27*). Remarkably, we observed that as thymic DC1s become CAT-experienced, they begin to express *Apoe* (Fig. 3F, left panel) in a similar manner to Claudin 1 in DC1s (Fig. 1E, 2F). As DC1 lineage maturation continued, strong upregulation of *Abcg1* and *Apol7c* genes that encode cholesterol efflux-associated proteins was detected (Fig. 3F, right panel). In regards to the *Nr1h2* gene, which is essential for the homeostatic maturation of splenic DC1s (*27*), we observed its highest upregulation in aDC1b among thymic immature and mature DC1 lineage subsets (Fig. 3F, right panel). In addition, our scRNAseq data showed that, in contrast to other thymic DC1 subsets, aDC1b expresses the marker, *Esam* (Fig. S5I), which has been associated with cholesterol efflux-dependent homeostatic mature splenic DC1s (*27*). It is of note that as thymic DC1s matured, distinct sets of genes that are associated with either homeostatic, both homeostatic and immunogenic but not immunogenic maturation alone, were gradually upregulated (Fig. S5J) (*27*, *35*). Taken together, our data points to a shared commonality in the process of thymic and splenic maturation of the DC1 lineage suggesting that in the thymus CAT is the driving force for homeostatic maturation.

### Claudin 1 facilitates positioning and contact between DC1s and mTECs

Given the interaction between CLAUDIN 1 on DC1s and CLAUDIN 3 on TRA-enriched mTEC subsets, we tested if mutual mTEC-DC1 positioning would be affected in the absence of Claudin 1. To visualize and measure the distance between mTECs and DC1s, we prepared two BM mixes that reconstituted a sub-lethally irradiated *Adig^GFP^* mouse, in which GFP marked Aire-expressing mTECs. These mixes, as depicted in Fig. 4A, gave rise to WT and Claudin 1-deficient DC1s at a 1:1 ratio, in which either WT (BM mix a) or Claudin 1-deficient (BM mix b) DC1s were labelled with TdTOMATO allowing the measurement of their distance from GFP^+^ mTECs (Fig. 4A-B).

**Figure 4.**
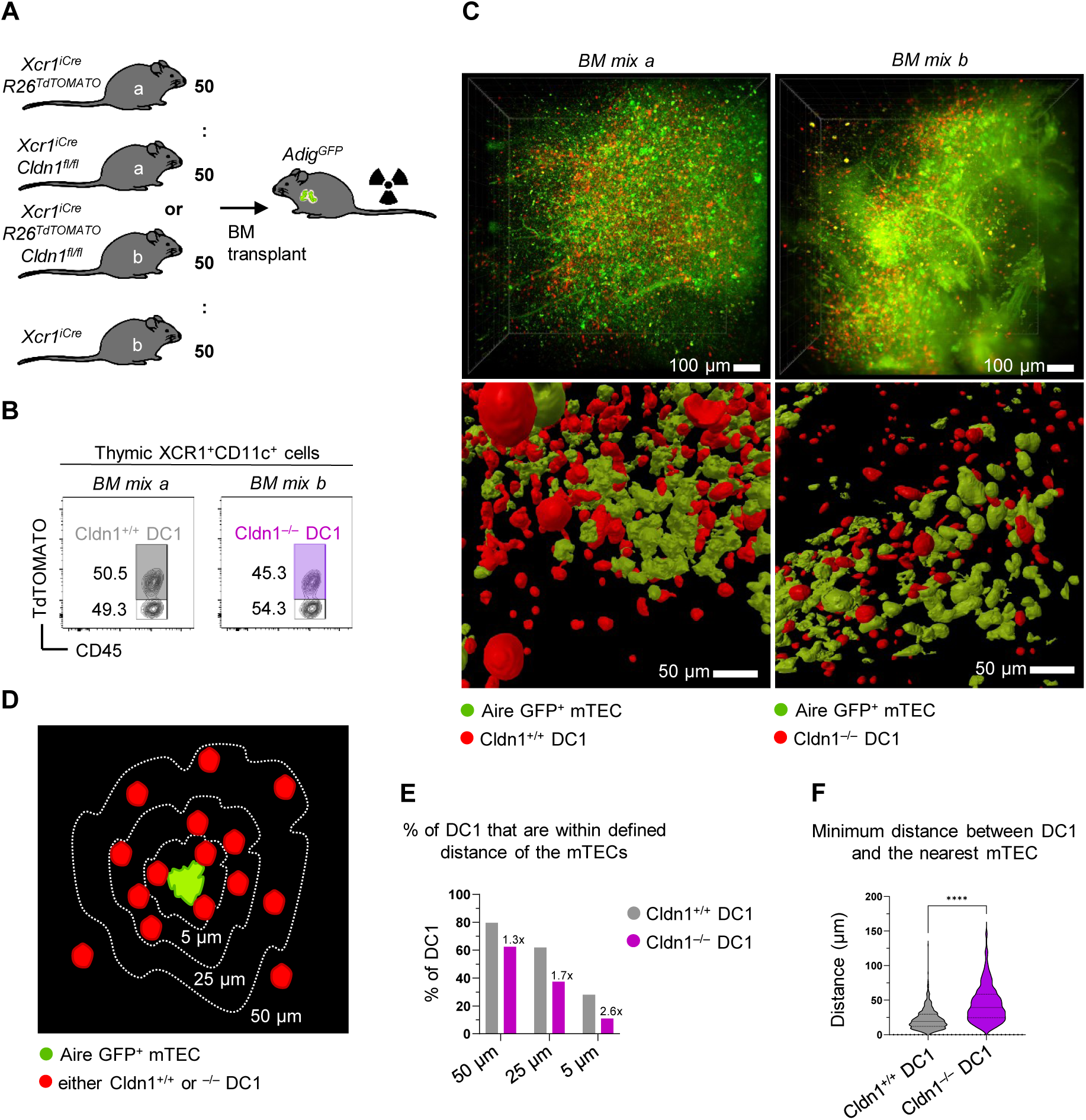
Claudin 1 facilitates positioning and contact between DC1s and mTECs. (**A**) Schematic of competitive BM chimera used to assess the role of Claudin 1 in positioning of DC1 lineage cells in the proximity of mTECs. Mouse models used as donors of BM are marked by the letters “a” and “b”. Note that *Adig^GFP^* recipients obtained BM either from mice “a” (BM mix a) or “b” (BM mix b). (**B**) Flow cytometry gating strategy used to analyze the reconstitution of competitive BM chimeras from Fig. 4A. Thymic CD11c^+^XCR1^+^ cells (DC1s) were gated as in Fig. S7A and either as TdTOMATO^+^ or TdTOMATO^−^. Note that TdTOMATO^+^ DC1s are Cldn1^+/+^ and TdTOMATO^−^ DC1s are Cldn1^−/–^ in the case of mice receiving BM mix “a” and TdTOMATO^+^ DC1s are Cldn1^−/–^ and TdTOMATO^−^ DC1s are Cldn1^+/+^ in the case of mice receiving BM mix “b”. (**C**) Light sheet fluorescence microscopy images of analogous regions within the thymic medulla of competitive BM chimeras from Fig. 4A-B. The top images capture the entire medullary compartment imaged. The bottom images visualize segmented objects (red DC1s and green mTEC clusters) within selected regions of the whole 3D images shown above. Separate legends are shown for BM mix “a” and “b”. (**D**) Schematic of the analysis of the regions imaged in Fig. 4C. The imaged area of each mTEC cluster captured was expanded by 5, 25, or 50 μm and the percentage of DC1s from the total within these expanded clusters was counted. Note that DC1s localized up to 5 μm from mTEC clusters are considered to be in direct contact with mTECs. (**E**) Percentage of DC1s that are within a defined distance of mTEC clusters related to Fig. 4D. The number above the Cldn1^−/–^ DC1 columns indicates the fold change reduction in the percentage of Cldn1^−/–^ DC1 with respect to the percentage of Cldn1^+/+^ DC1. (**F**) Violin plots showing minimum distance (μm) between DC1 and the nearest mTEC cluster related to Fig. 4C. Medians and quartiles are shown (n=2259 Cldn1^+/+^ DC1s and 531 Cldn1^−/–^ DC1s per representative experiment from a total of two experiments). Statistical analysis in F was performed using unpaired, two-tailed Student’s t-test, p < 0.0001 = ****.

Using a novel approach, we imaged the thymi of these competitive BM chimeras via light sheet fluorescence microscopy in combination with CUBIC clearance of thymic tissue (*51*), which allowed the imaging of large 3D regions of the thymic medulla (Fig. 4C, top panel). When we compared the thymic medulla of mice that received either BM mix a or b, we observed that in mix a, TdTOMATO^+^ DC1s (Cldn1^+/+^) were in close contact with the mTEC network, whereas in mix b, despite their proximity, TdTOMATO^+^ DC1s (Cldn1^−/–^) seemed to not directly contact mTECs (Fig. 4C, bottom panel). To quantify proximity, we determined the percentage of TdTOMATO^+^ DC1s (out of all TdTOMATO^+^ DC1s imaged) that were located within either 5, 25, or 50 μm from mTEC clusters (Fig. 4D). We observed that in general, the difference in the percentages within a given perimeter decreased as the distance increased (Fig. 4E). Importantly, in the case of DC1s located within 5 μm of the mTEC clusters, the percentage of WT DC1s (28%) was 2.6 times higher than Claudin 1-deficient DC1s (11%). Given that the 5 μm distance between cells was conducive for direct cell contact and a distance greater than 15 μm significantly reduced the likelihood of contact (*52*, *53*), our data suggests that Claudin 1 deficiency led to the dislocation of DC1s from mTECs, potentially compromising their cooperative characteristics in the establishment of central tolerance (*54*). In fact, the minimum distance between Cldn1^−/–^ DC1s and the nearest mTEC cluster was significantly longer compared to that of Cldn1^+/+^ DC1s (Fig. 4F).

### Claudin 1 in DC1 lineage regulates central tolerance

Although it has been shown that the DC1 cells are important for Treg selection (*16*, *22*), whether this function requires MHCII presentation has not been addressed. To this end, we used *XCR1^iCre^I-ab^fl/fl^* mice in which MHCII was ablated in the DC1 lineage (Fig. S6A-B) to analyze the Treg compartment (Fig. S6C). While the frequency of Foxp3^+^ Treg precursors (*55*) remained consistent, the frequency of Tregs, as well as CD25^+^ Treg precursors, was reduced (Fig. S6D). This strongly suggests that antigen presentation by the DC1 lineage governed the development of Tregs from their CD25^+^ but not Foxp3^+^ precursors. Surprisingly, the frequencies of both newly generated CD73^−^ as well as recirculating CD73^+^ Tregs were also significantly diminished (Fig. S6E) correlating with the marked reduction of splenic Tregs (Fig. S6F-G) (*34*). Hence, antigen presentation by the DC1 lineage is a crucial contributor to the generation of Tregs within the thymus.

Given the preferential pairing of DC1 subsets with TRA-expressing mTEC subsets (*10*), we analyzed the function of Claudin 1 in the DC1 lineage in relation to the selection of TRA-specific T-cells. We designed a novel *Defa6^iCre^R26^TdT-OVA^* mouse model (Fig. 5A) which contained a TdTOMATO-OVALBUMIN peptide-encoding fusion gene construct (TdT-OVA) within the R26 locus preceded by a STOP cassette flanked by loxP sites. This construct allowed the Cre-driven cell type-specific expression of TdT-OVA fusion protein. The OVA fragment is comprised of two peptides that are specifically recognized by TCR transgenic OTI and OTII T-cells (OTI/II cells). Since this Cre-driven TdT-OVA system mimics the expression of the Aire-dependent TRA, α-Defensin 6, and is recognized by OTII cells, it is suitable for the study of the generation of OVA-specific thymic Tregs.

**Figure 5.**
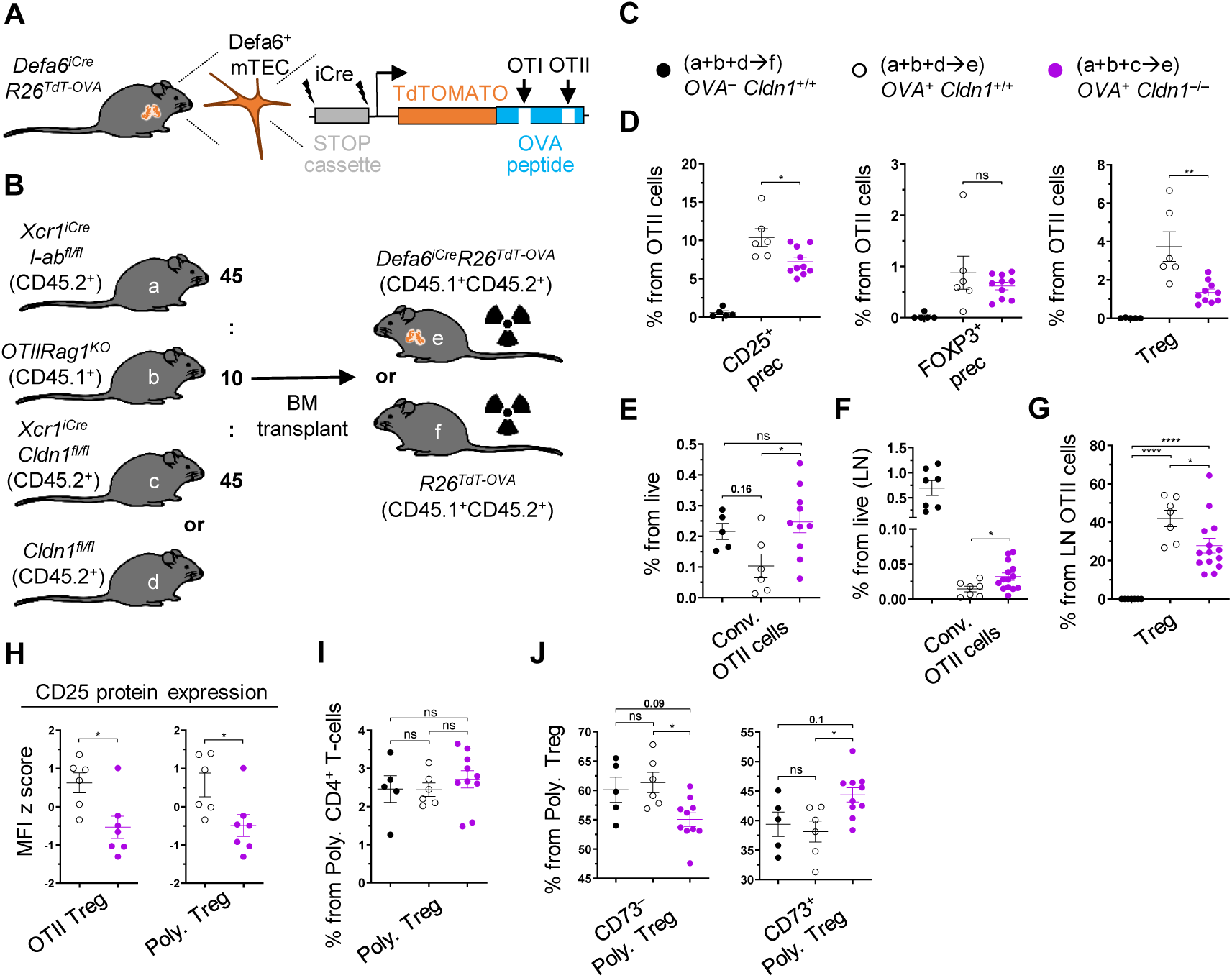
Claudin 1 in DC1 lineage regulates central tolerance. (**A**) Schematic of the generated *Defa6^iCre^R26^TdT-OVA^* model. (**B**) Schematic of donors and recipient genotypes used for competitive BM chimera experiments which assess the role of Claudin 1 in Treg selection and clonal deletion. Mouse models used are marked by the letters (a-f). Ratio for the preparation of BM mixtures is indicated. (**C**) Symbols used across Fig. 5, Fig. 6 and their supplements for the negative control samples (a+b+d→f, solid black circle), positive control samples (a+b+d→e, empty black circle) or test samples (a+b+c→e, violet circle) are shown. (**D**) Frequency of OTII Tregs and their CD25^+^ and Foxp3^+^ precursors within all OTII cells from thymi of competitive BM chimeras described in Fig. 5B-C (mean ± SEM, *n*=5-10 mice from 2 independent experiments). (**E**) Frequency of conventional OTII cells within all live cells from thymi of competitive BM chimeras described in Fig. 5B-C (mean ± SEM, *n*=5-10 mice from 2 independent experiments). (**F**) Frequency of OTII Tregs within OTII cells from skin-draining lymph nodes of competitive BM chimeras described in Fig. 5B-C (mean ± SEM, *n*=7-14 mice from 3 independent experiments). (**G**) Frequency of conventional OTII cells within live cells from skin-draining lymph nodes of competitive BM chimeras described in Fig. 5B-C (mean ± SEM, *n*=7-14 mice from 3 independent experiments). (**H**) MFI z score of CD25 expression within OTII Tregs and Polyclonal Tregs, the latter gated as in Fig. S7G, from BM chimeras described in Fig. 5B-C (mean ± SEM, *n*=6-7 mice from 2 independent experiments). (**I-J**) Frequency of Polyclonal Tregs from Polyclonal CD4^+^ T-cells (I) and newly generated (CD73^−^) and recirculating Tregs (CD73^+^) from Polyclonal Tregs (J) gated as in Fig. S7G from thymi of competitive BM chimeras described in Fig. 5B-C (mean ± SEM, *n*=5-10 mice from 2 independent experiments). Statistical analysis in D, F and H was performed using unpaired, two-tailed Student’s t-test and in E, G, I and J using one-way ANOVA with Tukeýs multiple comparisons, p ≤ 0.05 = *, p ≤ 0.01 = **, p ≤ 0.001***, p < 0.0001 = ****, ns = not significant.

To assess the effect of Claudin 1 deficiency in the DC1 lineage on the thymic selection processes of OTII cells, we used mixed BM chimeras in which 10% of the BM was of *OTIIRag1^KO^* origin (Fig. 5B, S7A-B). In this manner, we preserved the polyclonal T-cell repertoire to ensure normal development of the thymus (*56*, *57*) while keeping the OTII frequency low to provide optimal conditions for their conversion to Tregs (*58*). Furthermore, to introduce competition between Claudin 1-deficient and -sufficient DC1 lineages for OVA acquisition, in our test mouse model (*OVA^+^ Cldn1^−/–^*), we used a BM mixture composed of Cldn1^+/+^MHCII^−/–^ (from the *XCR1^iCre^I-ab^fl/fl^*mouse model) and Cldn1^−/–^MHCII^+/+^ DC1 (from the *XCR1^iCre^Cldn1^fl/fl^* mouse model) at a 1:1 ratio (Fig. 5B-C, S7A, S7C). Presumably, since Claudin 1 is required for close proximity of DC1s to TRA-loaded mTECs (Fig. 4, S4), Cldn1^−/–^MHCII^+/+^ DC1 would have limited access to mTECs with OVA in this setting and be positionally outcompeted by Cldn1^+/+^MHCII^−/–^ DC1. Conversely, even though Cldn1^+/+^MHCII^−/–^ DC1 would have access to OVA-producing mTECs, due MHCII deficiency, these cells will not be able to present OVA to OTII cells. However, since only the DC1 cells that can present antigens are those that lack Claudin 1, this experimental design can shed light on the impact of Claudin 1 deficiency on the indirect presentation of mTEC-derived antigens (*3*). Additionally, because Claudin 1 regulates DC1 maturation (Fig. 3, S5), the OVA presentation by Cldn1^−/–^MHCII^+/+^ DC1 lineage may be perturbed as well. It is of note that this defective selection does not only apply to OTII cells but all polyclonal T cells, which represent a majority of T cells in this BM-chimeric system. As a reference, we used reconstituted irradiated mice with a BM mixture from MHCII-deficient (Cldn1^+/+^MHCII^−/–^) and Claudin 1-sufficient WT mice (Cldn1^+/+^MHCII^+/+^) on OVA^−^ (*OVA^−^ Cldn1^+/+^*, negative control) or OVA^+^ (*OVA^+^ Cldn1^+/+^,* positive control) background (Fig. 5B-C). For flow cytometry analysis, T-cell subsets were gated as depicted in Fig. S7D.

We detected significantly decreased frequencies of OTII Tregs as well as OTII CD25^+^ Treg precursors in mice carrying the Claudin 1-deficient (*OVA^+^ Cldn1^−/–^*) DC1 lineage in comparison to the *OVA^+^ Cldn1^+/+^* chimeras (Fig. 5D). It should be noted that the *OVA^−^ Cldn1^+/+^* chimeras did not yield OTII Tregs or their precursors. This was expected since Treg generation requires presentation of their cognate antigen in the thymus (*59*). Analogous to the experiments with *XCR1^iCre^I-ab^fl/fl^* mice (Fig. S6E), the frequencies of OTII Foxp3^+^ Treg precursors were comparable between *OVA^+^ Cldn1^+/+^* and *OVA^+^ Cldn1^−/–^*chimeras (Fig. 5D). In addition, the frequencies of both newly generated CD73^−^ and recirculating CD73^+^ OTII Tregs were reduced (Fig. S7E) as in *XCR1^iCre^I-ab^fl/fl^*mice (Fig. S6F), indicating that defects in central tolerance were projected to the immune periphery. We also observed a slight increase in the frequency of conventional CD25^−^ Foxp3^−^ OTII cells in the thymi of *OVA^+^ Cldn1^−/–^* chimeras in comparison to *OVA^+^ Cldn1^+/+^* chimeras, suggesting incomplete clonal deletion (Fig. 5E). Since Claudin 1 deficiency affected both Treg selection and clonal deletion, we analyzed their impact on OTII cells in skin draining lymph nodes (Fig. S7F). Indeed, we detected a higher frequency of conventional OTII cells in *OVA^+^ Cldn1^−/–^* chimeras (Fig. 5F) and a reduction in the frequency of lymph node Tregs in comparison to the *OVA^+^ Cldn1^+/+^* chimeras (Fig. 5G), suggesting that both dominant and recessive central tolerance mechanisms were impaired in the absence of Claudin 1.

Interestingly, CD25 MFI in OTII Tregs was reduced in thymi of *OVA^+^ Cldn1^−/–^* mice when compared to the *OVA^+^ Cldn1^+/+^*control (Fig. 5H), implying that not only Treg frequencies but also their function may be compromised in the absence of Claudin 1 (*19*). Since the majority of the T-cell repertoire of our chimeras was polyclonal (Fig. 5B), we analyzed CD25 MFI on polyclonal Tregs and obtained a result similar to what was observed in OTII Tregs (Fig. 5H, S7G). Considering that the function of polyclonal Tregs may be affected as well, this prompted us to analyze Treg frequencies from developing polyclonal CD4^+^ T-cells (Fig. 5I). While these frequencies were comparable among all chimera variants analyzed, *OVA^+^ Cldn1^−/–^* mice exhibited a reduced frequency of newly generated Tregs in the thymus that was compensated by an increased frequency of recirculating Tregs (Fig. 5J). Taken together, this data suggests that Claudin 1 regulates the parameters of the thymic DC1 lineage that are critical for its tolerogenic functions and acts as an essential component of the molecular machinery of central tolerance impacting clonal deletion and Treg selection of TRA-specific T-cells.

### Claudin 1 deficiency in the DC1 lineage leads to break in tolerance and premature death

Since Claudin 1 is required for the induction of central tolerance, we investigated if its absence in DC1s would lead to autoimmunity. To this end, we collected sera from competitive BM chimeric mice (Fig. 5B-C) 30-40 weeks after transplantation and analyzed the presence of autoantibodies against kidney, liver, and stomach on composite tissue slides (Fig. 6A). Young WT mice and old *Aire^-/-^*mice (both non-irradiated) were used as negative and positive controls, respectively. In all three chimeric variants, we detected reproducible production of autoantibodies in comparison to the negative control (Fig. 6A-B) which is likely a consequence of mouse irradiation (*60*) or implementation of the *XCR1^iCre^I-ab^fl/fl^* mouse model (Fig. S6). However, while the serum autoantibody levels of the Claudin 1-sufficient chimeras (*OVA^−^ Cldn1^+/+^* and *OVA^+^ Cldn1^+/+^*) were comparable to the positive control (old *Aire^−/–^* mice) across the entire composite tissue images as well as in individual organs, *OVA^+^ Cldn1^−/–^* chimeras showed the highest autoantibody levels in all tissues analyzed (Fig. 6A-B). Surprisingly, autoantibody levels were similar between *OVA^−^ Cldn1^+/+^* and *OVA^+^ Cldn1^+/+^* chimeras, suggesting that autoantibody production was not triggered by self-reactive OTII T-cell clones but by polyclonal T cells. Indeed, while conventional OTII cells and OTII Tregs were still present in the immune periphery up to 40 weeks after BM transplantation (Fig. S8A-C), they were not enriched in the effector CD62L^−^CD44^+^ phenotype (*61*) in the mesenteric lymph node or in the spleen in *OVA^+^ Cldn1^−/–^* chimeras, suggesting that they were not responsible for the development of their autoimmune phenotype (Fig. S8D). Also, Paneth cells (PCs) as the only cells of the immune periphery with an active *Defa6* promotor (drives the production of OVA in *OVA^+^ Cldn1^+/+^* and *OVA^+^ Cldn1^−/–^*chimeras), showed similar numbers in the ileum of all BM chimeras (Fig. S8E-F), further supporting the negligible role of OTIIs in the observed autoimmunity.

**Figure 6.**
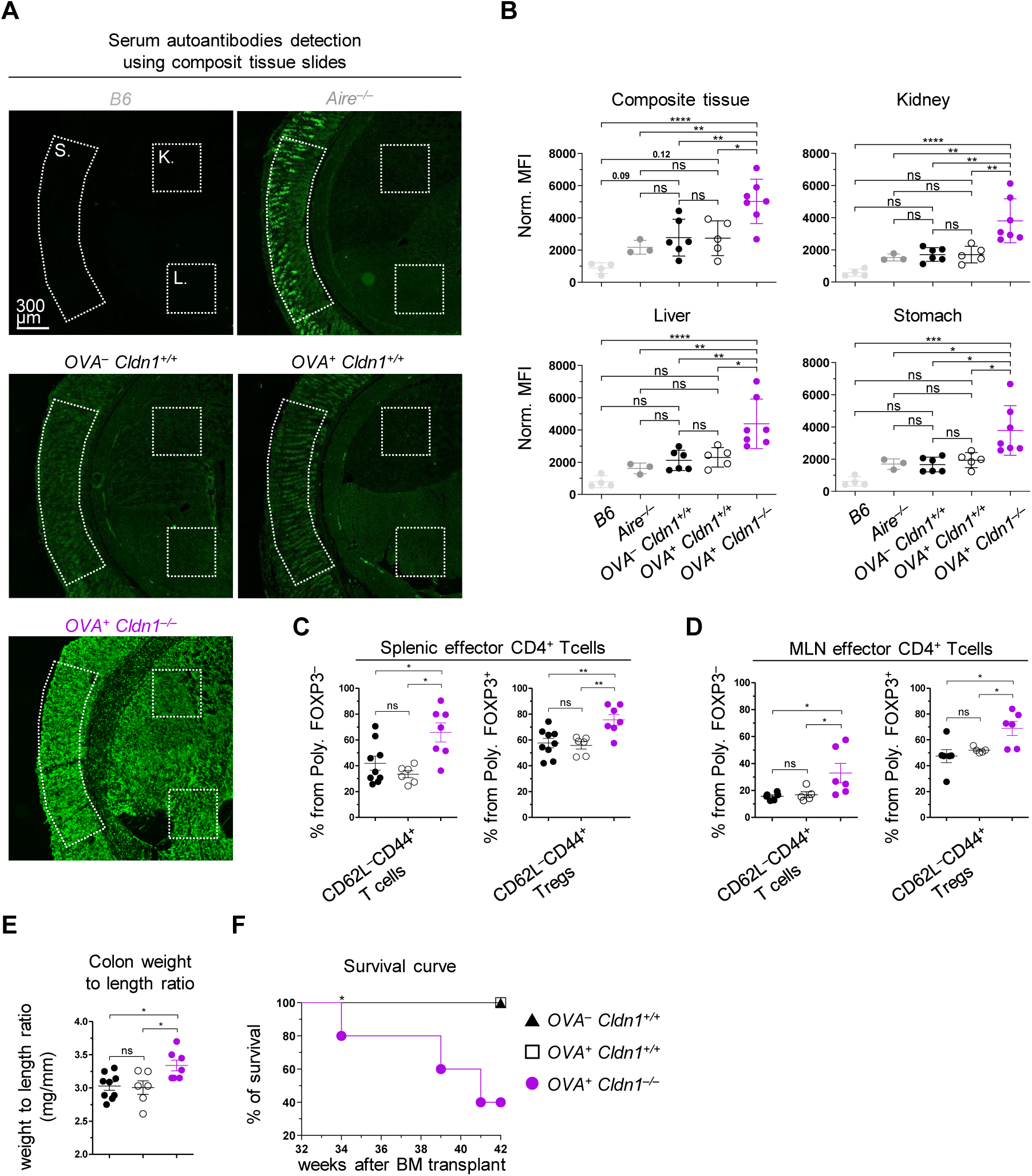
Claudin 1 deficiency in the DC1 lineage leads to break in tolerance and premature death. (**A**) Detection of autoantibodies from sera of BM chimeras from Fig. 5B-C using composite tissue slides. Serum autoantibody levels were quantified by the mean fluorescence intensity (MFI) of the secondary anti-mouse antibody conjugated to Alexa 555 (shown in green) across the entire image and from the regions demarcated by dotted lines corresponding to the individual tissues (K.= kidney, L.= liver, and S.= stomach) that make up the composite slide. Six week old *B6* WT and 30 week old *Aire^−/–^* mice on *BALB/c* background were used as negative and positive controls, respectively. Note that at the time of serum harvest, the competitive BM chimeras were 30-40 weeks post BM transfer. (**B**) MFI of serum autoantibody detection within the entire image of the composite tissue and individual tissues from Fig. 6A (mean ± SEM, *n*=3-7 mice from 2 independent experiments). (**C-D**) Frequency of effector memory CD62L^−^CD44^+^ cells within polyclonal conventional CD4^+^ T cells (Poly. FOXP3^−^) and Tregs (Poly. FOXP3^+^) gated as in Fig. S8A, isolated from mesenteric lymph nodes (MLN) (C) and spleens (D) of BM chimeras from Fig. 5B-C around 40 weeks post BM transfer (*n*=5-9 mice from 2 independent experiments). (**E**) Weight to length ratio of colons isolated from BM chimeras from Fig. 5B-C around 40 weeks post BM transfer (*n*=6-9 mice from 2 independent experiments). (**F**) Survival curve of BM chimeras from Fig. 5B-C (*n*=5 mice from 2 independent experiments). Color code as in Fig. 5C. Statistical analysis in B, C, D and E was performed using one-way ANOVA with Tukeýs multiple comparisons test and in F using Log-rank (Mantel-Cox) test, p ≤ 0.05 = *, p ≤ 0.01 = **, p ≤ 0.001***, p < 0.0001 = ****, ns = not significant.

On the other hand, the polyclonal CD4^+^ T cells, showed an increase in the effector CD62L^−^CD44^+^ phenotype in the conventional compartment of spleens of *OVA^+^ Cldn1^−/–^* chimeras compared to controls (Fig. 6C, S8A). However, we observed an even more significant increase of these cells in the Treg compartment which together reflected systemic inflammation that correlated with a break in tolerance. Moreover, we detected the same trend within polyclonal CD4^+^ T cells from mesenteric lymph nodes of *OVA^+^ Cldn1^−/–^* chimeras (Fig. 6D, S8A), indicating inflammation in the intestines. In fact, compared to controls, we detected a significant increase in the colon weight to length ratio in the *OVA^+^ Cldn1^−/–^* chimeras (Fig. 6E, S8G), which is a telltale sign of colitis.

Remarkably, this systemic break in tolerance correlated with the premature death of Claudin 1-deficient chimeras (*OVA^+^ Cldn1^−/–^*) (Fig. 6F). Specifically, while all *OVA^−^ Cldn1^+/+^* and *OVA^+^ Cldn1^+/+^*chimeric mice were still alive 42 weeks after BM transplantation, 60% of the *OVA^+^ Cldn1^−/–^* chimeras did not survive beyond this time point. Hence, the ablation of Claudin 1 in the DC1 lineage resulted in the shortened lifespan of the mice which was accompanied by a break in immune tolerance.

## DISCUSSION

In this study, we identified the tight junction protein, CLAUDIN 1, which plays a substantial role in antigen transfer-coupled maturation of the thymic DC1 lineage. We annotated the phenotypic heterogeneity of this lineage and performed tracing experiments that confirmed that immature DC1s transition to early mature aDC1a and subsequently to late mature aDC1b. A DC1 lineage-specific knockout of the *Cldn1* gene showed that Claudin 1 is critical for the maturation of aDC1a and aDC1b. Since the CAT-experienced DC1 lineage maturation was accompanied by the upregulation of cholesterol efflux associated genes, this suggested that CAT was responsible for the homeostatic maturation of DC1s. We detected the highest expression of CLAUDIN 3, the binding partner of CLAUDIN 1, on TRA-expressing mTECs, which lends credence to the phenomenon of their preferential pairing with XCR1^+^ and XCR1^−^ aDCs (*10*). The importance of Claudin 1 was confirmed by light sheet fluorescence microscopy, which illustrated that it participates in the establishment of central tolerance by ensuring optimal positioning of DC1s within the mTEC network. We also demonstrated the importance of MHCII expression on the DC1 lineage for Treg selection. Finally, we generated a novel *Defa6^iCre^R26^TdT-OVA^* mouse model with OVA neo-self-antigen that mimicked TRA expression and showed that Claudin 1 deficiency in the DC1 lineage diminished clonal deletion of OVA-specific T-cells as well as their selection into Tregs. Consistent with impaired central tolerance, we detected high serum titers of autoantibodies against several tissues, increased frequency of effector CD4^+^ T cells and Tregs, and symptoms of colitis. This break in tolerance correlated with the shortened lifespan of these animals. In aggregate, we uncovered a novel molecular mechanism that employed Claudin 1 as an essential molecule in CAT-coupled maturation of cells of the DC1 lineage, which in turn provides an antigen-presenting network for proper thymic selection of TRA-specific T-cells.

The compilation of our data consolidates the current knowledge regarding the classification of aDC subsets and their maturation path. The scRNAseq thymus atlas developed by *Park and colleagues* divided aDCs into two conventional DC lineages: *Xcr1^+^* aDC1 and *Sirpa^+^* aDC2, along with an aDC3 subset, which exhibited dramatically reduced or lack of expression of *Xcr1* and *Sirpa* as well as other lineage-specific markers (*62*). Recently, *Bosteels and colleagues* described two maturation states of splenic aDC1s, early and late, the latter that also showed decreased levels of *Xcr1* (*27*, *35*). Our scRNAseq confirmed that both the DC1 and DC2 lineages undergo maturation from early (aDC1/2a) to late stages (aDC1/2b). Along with maturation from the “a” to “b” state, accompanied by the gradual diminishment of *Sirpa* or loss of *Xcr1*, we unexpectedly observed upregulation of *Cd81 and Il7r* genes that are affiliated with DC1 and DC2 lineages, respectively. Therefore, these genes can be used as surrogate markers for the late mature stages, aDC1b and aDC2b, when *Xcr1* and *Sirpa* are absent and downregulated, respectively. In contrast to our lineage tracing experiments, attempts to track the development of either thymic or extrathymic DCs failed to detect XCR1^−^ aDC1b due to the gating of only XCR1^+^ DCs (*27*, *30*). Therefore, in the thymus, aDC1a represents early mature XCR1^+^ DC1s which in the past were referred to as mDC1, CCR7^+^ DC1, mregDC1, or aDC1, while aDC1b represents late mature XCR1^−^ DC1s (*10*, *18*, *26*, *27*, *30*, *32*, *62*).

We have shown for the first time that the mechanism that drives thymic DC1 homeostatic maturation is similar to what has been described in splenic DC1s (*27*, *30*). Surprisingly, homeostatic maturation of DCs, unlike immunogenic maturation, does not involve the engagement of pattern recognition receptors (*35*), yet it leads to nearly the same changes in the transcriptome of DCs regardless of the type of lymphoid tissue, i.e. thymus, spleen or lymph nodes (*30*). However, DC1 homeostatic maturation differs in transcriptomic changes related to genes involved in fat metabolism and interferon sensing (*27*, *30*, *32*). Consistent with this notion, we detected the upregulation of cholesterol efflux-associated genes in CAT-experienced DC1s, which continued along the maturation process. This result is in agreement with a scenario where mTEC-derived cholesterol-containing apoptotic bodies induce the liver X receptor pathway, driving the efflux of cholesterol which induces homeostatic maturation of the thymic DC1 lineage, analogous to what has been proposed in splenic DC1 maturation (*27*). In this context, a complementary thymus-specific mechanism that is responsible for DC1 maturation, which depends on type III interferon sensing has been recently reported (*32*). In contrast to DC1s, the DC2 lineage was found to be non-responsive to apoptotic cell engulfment in terms of its maturation (*27*). Instead, thymic DC2s display a type 2 cytokine gene expression signature requiring IL4R signalling to become mature and engage in clonal deletion (*18*). While the main drivers of maturation pathways are DC lineage specific, homeostatic maturation of both thymic DC1 and DC2 requires the presence of T-cells, TCR-MHC interactions, and to a lesser extent, CD40 signalling (*31*).

As alluded to in the results section, we have shown that Claudin 1 is required for juxtaposition of DC1s to mTECs within the epithelial network. This may represent an analogous scenario that has been observed in the gut or skin where DCs cross the epithelial barrier in a Claudin 1-dependent manner to capture and subsequently present antigens (*41*, *44*). Our observation of the highest expression of Claudin 3, which is the ligand of Claudin 1, on mTECs that produce Aire-dependent TRAs strongly supports this hypothesis. Interestingly, recently discovered mimetic cells (*14*, *15*), which express high levels of TRAs but may possess only a limited antigen presentation capability (*3*), also display high levels of Claudin 3. This, along with mTEC^HI^ cells, predisposes mimetic cells to be prime targets for the Claudin 1^+^ DC1 lineage to acquire TRAs via CAT. Other CAT-associated molecules identified in our scRNAseq screening fit within this framework including the integrin, CD103, (encoded by *Itgae*) which mediates adhesion of DCs to the epithelium (*63*). To accomplish CAT, molecules such as scavenger CD36 are then required for the acquisition of mTEC antigens. Considering that the execution of CAT is likely the result of cooperative action of several molecular determinants that are expressed on DC1s or by their mTEC interacting partners, we propose that the loss of function of any of these determinants will be compensated by other molecules. Strikingly, since the ablation of Claudin 1 in the DC1 lineage had no gross effect on the efficiency of CAT but severely impaired the frequency of TdTOMATO^+^ aDC1b, this suggests two possible scenarios: (i) in respect to CAT, the absence of Claudin 1 can be readily substituted by other determinants or (ii) Claudin 1 is necessary only for the CAT-coupled maturation of the DC1 lineage.

Unexpectedly, Claudin 1 deficiency resulted in the profound reduction of only TdTOMATO^+^ aDC1b, even though Claudin 1 expression was low in this subset. We presumed that Claudin 1 deficiency should affect aDC1a since this subset expressed the highest levels of Claudin 1. Even though we annotated aDC1a and aDC1b as separate subsets, the developmental process from aDC1a to aDC1b represents a continuum of the maturation state. Since Claudin 1-deficient aDC1a showed a higher frequency of TdTOMATO^+^ cells as opposed to its Claudin 1-sufficient counterpart, this seems to indicate that Claudin 1 controls maturation of CAT-experienced DC1s. Notably, the observation that Claudin 1 deficient DC1 lineage tends to accumulate TdTOMATO^+^ cells in the aDC1a state, which is reflective of its stalled development, suggests that Claudin 1 is critical for the transition from aDC1a to aDC1b. In addition, since the cellularity of Claudin 1-deficient TdTOMATO^+^ aDC1a compared to its Claudin 1-sufficient counterpart in the same BM competitive chimeric mice was decreased, it is not implausible that Claudin 1 influences not only maturation but also the survival rate of this subset. Regardless of the mechanism through which Claudin 1 regulates DC1 maturation, the most important finding of this study is that in a Claudin 1-deficient setting, the aDC1b subset was greatly impaired. Since aDC1b is a fully matured subset which specializes in antigen presentation, its dramatically reduced numbers explain how defective DC1 maturation leads to a break in central tolerance.

Interestingly, the role of Claudin 1 in DC1 maturation may be explained from previous findings that showed that the Claudin family of proteins interacts with several classes of non-tight junction molecules such as tetraspanins (*64*). It is of particular interest that Claudin 1 interacts (on the same cell) with the tetraspanin family member, CD81, which is a molecular sensor of cholesterol (*65*, *66*). As we previously noted, cholesterol sensing during apoptotic cell acquisition drives homeostatic maturation of the DC1 lineage (*27*). Our finding showing the gradual expression of *Cd81* during the DC1 maturation suggests that Claudin 1 may drive the maturation of CAT-experienced DC1 through its interaction with CD81, which is linked to cholesterol sensing.

Taking advantage of our *Defa6^iCre^R26^TdT-OVA^* mouse model, we have shown that in the absence of Claudin 1 in the thymic DC1 lineage, the immune periphery exhibited an elevated frequency of self-reactive conventional T-cells and a reduced frequency of Tregs, attesting to the leakiness of central tolerance. In fact, *Defa6^iCre^R26^TdT-OVA^* chimeric mice carrying the Claudin 1 deficient DC1s manifested an autoimmune phenotype which closely mimicked the symptoms of other mouse models that suffer from insufficient presentation of TRAs in the thymus, such as *Aire^−/–^* mice, which are characterized by high autoantibody titers, multi-organ autoimmunity, and shortened life expectancy (*67*, *68*). In general, dysregulation of Claudin 1 expression contributes to numerous autoimmune diseases including rheumatoid arthritis, type 1 diabetes, multiple sclerosis, or inflammatory bowel disease (*69–72*). While *Cldn1^-/-^* mice die within the first day of life due to trans-epidermal water loss (*73*), the Claudin 1 knockdown showed a disintegration of the epidermis that resembled atopic dermatitis (*33*). Thus, the consensus is that abnormal Claudin 1 expression in the epithelium lead to the loss of barrier integrity and subsequent breakdown of peripheral tolerance. In this study, we proposed a different mechanism by which Claudin 1 deficiency leads to autoimmunity by the loss of aDC1b which limits the ability of central tolerance to delete self-reactive T-cells or convert them into Tregs (*16*, *59*). This work further extends the understanding of the importance of the DC1 lineage for antigen presentation in peripheral tolerance as previously described (*34*) to the mechanisms of central tolerance described in this study. Collectively, Claudin 1 deficiency leads to autoimmunity indirectly through the loss of the epithelial barrier integrity or directly through the failure of the DC1 lineage to mature and to purge self-reactive T-cells.

## MATERIALS AND METHODS

### Study design

The overarching aim of this study was to search for molecules involved in cooperative antigen transfer (CAT) and/or CAT-coupled processes, i.e. DC maturation, in the DC1 lineage using scRNAseq, flow cytometry, and competitive BM chimeras (mice) that allowed tracking of CAT. We focused our study on the tight junction protein CLAUDIN 1. Using a Claudin 1 conditional knockout mouse in the DC1 lineage, we established its relevance in CAT-coupled maturation of thymic DC1s. To address this, we first re-analyzed the heterogeneity of DC1 maturation states using scRNAseq, a reporter mouse, and BrdU lineage tracing and discovered previously overlooked late mature DC1s: aDC1b. We hypothesized that Claudin 1 may be involved in DC1 biology by enabling their optimal positioning in the vicinity of mTECs and addressed this using light sheet fluorescence microscopy in combination with the CUBIC clearing protocol. We generated a novel mouse model, *Defa6^iCre^R26^TdT-OVA^*, to test whether Claudin 1 deficiency negatively affects the selection of TRA-specific T cells in the thymus. We also used this model to determine whether Claudin 1 deficiency in DC1s translates into symptoms of autoimmunity. Light sheet fluorescence microscopy, autoantibody detection, and Paneth cell counting experiments were imaged and analyzed blinded by a source unfamiliar with the genotype/phenotype of the mice. Mice and tissue samples were excluded from the analysis if BM reconstitution was insufficient or cell isolation was suboptimal.

### Mice

All mice used in this study were on the full *C57Bl/6J (B6)* background, except for *Aire^−/–^* mice which were on full *BALB/c* background, and bred under SPF conditions at the animal facility of the Institute of Molecular Genetics of the Czech Academy of Sciences (IMG) and University of Birmingham, Biomedical Services Unit, UK. Experimental procedures with mice were approved by the ethical committee of the IMG, Birmingham Animal Welfare and Ethical Review Board and UK Home Office. Mice were fed by standard rodent high-energy diet and given reverse osmosis-filtered water ad libitum. Mice were bred under light/dark cycle that oscillated every 12 hours and in constant temperature and humidity of 22 ± 1°C and 55 ± 5%, respectively. *B6*, *Foxn1^Cre^* (B6(Cg)-*Foxn1^tm3(cre)Nrm^*/J; #018448) (*74*), *Ly5.1* (B6.SJL-*Ptprc^a^ Pepc^b^*/BoyJ; #002014) (*75*), *I-Ab^fl/fl^* (B6.129X1-*H2-Ab1^b-tm1Koni/^*J; #013181) (*76*), *Rag1^−/−^* (B6.129S7-*Rag1^tm1-Mom^*/J; #002216) (*77*), and OT-II (B6.Cg-Tg(TcraTcrb)425Cbn/J; #004194) (*78*) mice were purchased from the Jackson Laboratories. *R26^TdTOMATO^* mice (B6;129S6-*Gt(ROSA)26Sor^tm14(CAG-tdTomato)Hze^*/J; #007908) (*47*) were provided by V. Kořínek (IMG, Prague, Czech Republic). *Defa6^iCre^* mice (*79*) were kindly provided by R. S. Blumberg (Division of Gastroenterology, Department of Medicine, Brigham and Women’s Hospital, Harvard Medical School, Boston, MA). *XCR1^iCre^* mice (*34*) were kindly provided by B. Malissen (Centre d’Immunologie de Marseille-Luminy, Aix Marseille Université, Inserm, CNRS, Marseille, France). *Cldn1^fl/fl^*mice (*33*) were kindly provided by S. Tsukita (Advanced Comprehensive Research Organization, Teikyo University, Tokyo, Japan). *Adig^GFP^* (*80*) and *Aire^−/–^* (*81*) mice were kindly provided by L. Klein (Ludwig Maximilian University, Munich, Germany). To harvest murine tissues, mice were euthanized by cervical dislocation at 4-7 weeks of age, except for bone marrow (BM) chimera experiments, where BM was transplanted into sub-lethally irradiated mice at 5-8 weeks of age. These mice were euthanized 5-6 weeks after BM transplantation, except for the mice subjected for the analysis of autoimmunity symptoms, which were culled 30-40 weeks after BM transplantation or used for the survival curve experimentation. In addition, *Aire^−/–^* mice were euthanized at 30 weeks of age. In all individual experiments, littermates were used regardless of their sex and caging. BM donors and mice used for scRNAseq were females.

### *R26^TdT-OVA^* mouse model

To generate mice with inducible TdTOMATO-OVA (TdT-OVA) expression, we designed and synthesized a plasmid vector (GenScript) for site-specific integration into a mouse Rosa26 (R26) locus. This vector includes a CAG promoter, a loxP-STOP-loxP cassette, the TdT-OVA transgene, a Woodchuck hepatitis virus posttranscriptional regulatory element (WPRE), and a bovine growth hormone polyadenylation signal (bGHpA). We flanked the knock-in sequence with R26 homology arms (*82*) and gRNA target sequences (5‘-CTCCAGTCTTTCTAGAAGATGGG-3’), to facilitate the efficiency of site-specific integration (*83*). Using the online software CRISPOR Design Tool ([http://crispor.tefor.net/]), we designed a R26 targeting gRNA (5‘-CTCCAGTCTTTCTAGAAGAT-3’). For pronuclear microinjections, we combined the targeting vector with gRNA and Cas9 protein and then introduced them into C57Bl6n-derived zygotes as previously described (*82*). Founder animals carrying site-specific insertion were identified using PCR with primers flanking homology arms. The full-length knock-in sequence was verified by Sanger sequencing. The expression of TdT-OVA protein by *R26^TdT-OVA^*mice crossed to *Itgax^Cre^* (*84*), *Foxn1^Cre^* or *Defa6^iCre^* mice was verified by flow cytometry. Specifically, we tested the presence of OTII peptide in Fig. 5 and S7, OTI peptide by SIINFEKL staining and proliferation assay of OTI TCRtg T-cells and TdTOMATO expression and its CAT by crossing *R26^TdT-OVA^* mice to *Foxn1^Cre^* and *Defa6^iCre^* strains.

### Cell isolations

Thymi were excavated using forceps, cut into 10-15 pieces and digested with an enzymatic cocktail of 0.1 mg/ml Collagenase D (Roche) and DNAse I (40 U/ml; Roche) dissolved in RPMI 3% FBS medium. Note that pieces of each thymus were put into 1 ml of enzymatic cocktail in a 1.5 ml eppendorf tube. To isolate thymic epithelial cells, 0.1 mg/ml Dispase II (Gibco) was added to the enzymatic cocktail. To complete digestion, enzymatic cocktails containing thymi were put into thermoshaker and incubated for ∼80 minutes at 37°C while shaken at 800 rpm. After incubation, non-digested thymic pieces were pipetted up and down several times using a cut pipet tip until the solution was homogeneous and then filtered into a 15 ml falcon tube. To stop the enzymatic reaction, each thymic solution was washed with 2 ml of ice cold 3% FBS and 2 mM EDTA solution in PBS. Then, thymi were spun down (4°C, 400 x g, 10 min). In the case of T-cell isolation, pellets were resuspended in 1 ml of ACK lysis buffer, incubated for 3 minutes, washed with 14 ml of 3% FBS and 2 mM EDTA solution in PBS and spun down (4°C, 400 x g, 10 min). 1/10 of pellet was used for later analysis. In the case of thymic myeloid APCs or thymic epithelial cell isolations, Percoll (Cytiva) enrichment was conducted by resuspending the pellets in 2 ml PBS, underlaid with 2ml of 1.065 g/ml Percoll and then 2 ml of 1.115 g/ml Percoll to create three separate layers. Next, the samples underwent gradient centrifugation (4°C, 1500 x g, 30 min, w/o break and acceleration). After centrifugation, two cell layers were formed. The bottom layer consisted of smaller cells such as T-cells and erythrocytes and the upper layer of myeloid APCs and thymic epithelial cells. The cells from the upper layer were gently transferred into 10 ml of 3 % FBS and 2 mM EDTA solution in PBS and spun down (4°C, 300 x g, 10 min). The resulting pellet was used for further analysis. In the case of isolation of splenic T-cells or T-cells from lymph nodes, the same approach that was used for the isolation of thymic T-cells was applied. Note that 1/5 of the spleen was used for cell isolation and such 1/5 was cut into ∼10 pieces. Lymph nodes were opened using 26G needle. The requirements for the isolation of cells designated for scRNAseq are described in its dedicated paragraph.

### Flow cytometry analysis

To stain cell surface markers for flow cytometry (FACS) analysis, cells were incubated with antibodies or other staining reagents at 4°C in the dark for 20–30 min. Note that biotin conjugated antibodies were stained prior to staining with fluorochrome-conjugated streptavidin and antibodies against other surface markers at 4°C in the dark for 20–30 min. In the case of anti-CCR7 antibody (BioLegend) staining, incubation was conducted on a thermoshaker at 37°C, 800 rpm for a minimum of 30 min prior to all other staining incubations. After each incubation, cells were washed using 1 ml of 3% FBS and 2 mM EDTA solution in PBS and spun down (4°C, 300 x g, 10 min). Note that in the case of T-cell staining, the centrifugation force was 400 x g. To stain intracellular markers, cells were fixed using a Foxp3 staining kit (Thermofisher) for 30 min after surface staining according to manufactureŕs protocol. Fixed cells were incubated with primary antibodies for 30 min at room temperature (RT) and in the case of unconjugated primary antibodies staining, an additional 15 min staining at RT with secondary antibodies was conducted. After each incubation, cells were washed using 10x diluted Permeabilization buffer (Thermofisher) and spun down (4°C, 500 x g, 10 min). Dead cells were excluded using either Hoechst 33258 (Sigma-Aldrich) or fixable viability dye eFluor 506 (eBioscience). FACS analysis was performed using FACSymphony A5, LSRFortessa, FACSDiva software, and FlowJo V10 software (BD). The abbreviation MFI used in the figures and figure legends stands for the median fluorescence intensity. A list of antibodies and other staining reagents can be found in Table S1.

### Single-cell RNA sequencing

To perform scRNAseq of thymic myeloid APCs, thymi from 6-week-old *Foxn1^Cre^R26^TdTOMATO^* mice were enzymatically digested as described (see Cell isolations). Importantly, isolated cells were not subjected to Percoll enrichment but were resuspended in a cocktail of anti-CD11c and anti-CD11b antibodies both conjugated with biotin, stained on ice for 25 min, washed with 3 % FBS and 2 mM EDTA solution in PBS, and then spun down (4°C, 300 x g, 10 min). Next, cells were stained with anti-biotin magnetic beads (Miltenyi) according to manufactureŕs protocol and CD11c^+^ and CD11b^+^ cells were MACS-enriched using QuadroMACS (Miltenyi). Enriched cells were stained with fluorochrome-conjugated streptavidin on ice for 15 min, washed with 3 % FBS and 2 mM EDTA solution in PBS and then spun down (4°C, 300 x g, 10 min). Next, ∼ 1 x 10^5^ of Streptavidin^+^ cells were sorted using a BD Influx cell sorter (BD) from a pool of three female littermate thymi. Importantly, half of the cells were sorted as TdTOMATO^+^ (CAT-experienced cells) and the other half as TdTOMATO^−^ (CAT-inexperienced cells) into separate collection tubes to prepare two individual scRNAseq libraries (samples). To check that viability of sorted cells was greater than 90%, an automated TC20 cell counter (Bio-Rad) was used. ScRNAseq libraries were prepared using a Chromium controller and the Chromium Next Gen single-cell 3’ reagent kit version 3.1 (both 10X Genomics) according to the manufactureŕs protocol targeting 4000 cells per sample, i.e. 4000 of TdTOMATO^+^ as well as TdTOMATO^−^ cells sequenced. The quality and quantity of the resulting cDNA and libraries were determined using an Agilent 2100 Bioanalyzer (Agilent Technologies). The sample libraries were sequenced in a single run of NextSeq 500 instrument (Illumina) using a high-output kit with mRNA fragment read length of 56 bases. We used 10X Genomics Cell Ranger software suite (version 3.1.0) to quantify gene-level expression based on GRCm38 assembly (Ensembl annotation version 98) (*85*).

### Bioinformatic analysis

For bioinformatic analysis of 10x scRNAseq data, we used a standard Seurat (v 4.0.2) pipeline, which was performed in R v 4.0.2 (R core team 2020) in a similar setting as done previously (*86*). In brief, 10x Cell Ranger raw read counts were used as an input. Cells were filtered to obtain those containing at least 1000 detected features (genes) and less than 20% of mitochondrial RNA read counts. Samples were then pooled and processed together. Cell types were annotated using a combination of clustering and canonic cell type-specific marker genes expression profile. If a cluster contained a mixed population, sub-clustering was performed. Afterwards, the cell type definition dataset was filtered to obtain clusters containing only conventional dendritic cells and monocyte/macrophage lineages. This filtered dataset was again clustered and sub-lineage cell types were defined using the same method as noted above. Seurat embedded differential expression analysis was used to define genes whose expression accompanies CAT in individual cell types as well as additional cell type markers.

### Bone marrow chimeras

To prepare bone marrow (BM) chimeras, femurs and tibias of euthanized BM donors were isolated, cleaned of surrounding tissues, and cut at both ends. BM was flushed out from the bones using PBS and syringe with 26G needle into a 15 ml falcon tube. Isolated BM was spun down (4°C, 400 x g, 10 min). The obtained pellets were resuspended with 1 ml of ACK lysis buffer, incubated for 3 minutes, washed with 14 ml of 3% FBS and 2 mM EDTA solution in PBS, and spun down (4°C, 400 x g, 10 min). Recipient mice were sub-lethally irradiated with 6 Gy and each mouse received 2 × 10^6^ of isolated BM cells through the tail vein. Importantly, prior to the transplantation, BMs were mixed at a 50:50 (Fig. 3A, 4A) or 45:45:10 ratio (Fig. 5B). After transplantation, the mice were monitored on a daily basis for signs of infection/wasting following irradiation. To protect the mice against infection, we supplemented their water with 2ml/100 ml of gentamicin (Aagent) for two weeks. The efficiency of reconstitution of mixed BM chimeras is shown in Fig. 4B, S5F-G, and Fig. S7A-C.

### BrdU lineage tracing

To trace DC1 lineage, WT *Ly5.1* mice were injected with 1.5 mg of BrdU i.p. and culled at one, two, three, four, five or seven day(s) after BrdU administration. Thymi of culled mice were enzymatically digested and thymic DCs were isolated. To stain BrdU^+^ thymic DCs, an established protocol was used (*87*). Briefly, thymic DCs were resuspended in 100 μl of BD fix/perm (BD) and incubated for 30 min at 4°C. Next, cells were washed in 1ml Perm/wash (BD) and spun down (4°C, 500 x g, 10 min). The pellet obtained was resuspended in 100 μl of BD Cytoperm buffer Plus (BD), incubated for 10 min on ice and washed. Afterwards, another round of fixation in 100 μl of BD fix/perm for 5 min on ice was conducted followed by washing. Next, cells were treated with DNAse I (1 mg/ml) for 45 min at 37 °C and washed. Then, DNAse-treated cells were stained with anti-BrdU FITC antibody diluted 1:100 in 100 μl Perm/wash buffer for 20 min at RT. After staining, cells were washed and subjected to FACS analysis.

### Light sheet fluorescence microscopy

For the segmentation, distance calculation, and visualization of thymic DC1s and mTECs, Light sheet fluorescence microscopy was used. Thymi of *Adig^GFP^* chimeras possessing mixed BMs were harvested 6 weeks after BM transplantation. Thymic lobes were separated and one was used for flow cytometry analysis of BM reconstitution. Thymic lobes used for microscopy were first fixed overnight at 4 °C in 3.8% paraformaldehyde. After thorough washing, the samples were cleared using a modified Clear Unobstructed Brain Imaging Cocktail and Computational (CUBIC) protocol (*88*, *89*). In the first step, the samples were cleared for 5 days at 37 °C in CUBIC1 solution (35 wt% dH2O, 25 wt% urea, 25 wt% N,N,N’,N’-Tetrakis(2-Hydroxypropyl) ethylenediamine (4NTEA), 15 wt% Triton X-100). The cleared samples were rinsed for one hour in CUBIC wash solution (0.5% BSA, 0.01% sodium azide, 0.01% Triton X-100 in PBS) three times. Subsequently, the samples were incubated in CUBIC 2 clearing solution (23.4 wt% dH2O, 22.5 wt% urea, 9 wt% triethanolamine (TEA), 45 wt% sucrose, 0.1% (v/v) Triton X-100) for 3 days to achieve refractive index matching. Analogical medullary regions of the same size were then visualized using a Zeiss Z.1 Light Sheet Microscope equipped with a 1.45 RI clearing chamber. A total of three channels was acquired: green (excitation: 488 nm, detection: 498 nm) for GFP signal detection, red (excitation: 561 nm, detection: 571 nm) for TdTOMATO signal detection, and a cross-channel (excitation: 488 nm, detection: 571 nm) to capture autofluorescence. This autofluorescence helped to distinguish positive signals in the two specific channels during subsequent steps. The raw data was first 3D deconvolved using Huygens Professional software (*89*) and subsequently analyzed in Arivis 4D (version 4.2.) with the following steps. First, voxel training was performed for the machine learning segmenter using all three channels until the model reliably recognized DC1s and mTEC clusters from the background. Subsequently, the TdTOMATO^+^ DC1s and GFP^+^ mTEC clusters were segmented. In the second step, splitting was performed for DC1s with a sensitivity of 57.33%. In the third and fourth steps, both categories were filtered and all artifacts and fragments smaller than 400 μm^3^ were removed. Subsequently, the minimum distances between each DC1 and the nearest mTEC cluster were measured using the Distances module. Finally, three expansions of the mTEC clusters were performed sequentially in the Compartments module, by 5, 25 and 50 μm, and the percentage of DC1s from total in each of the expanded clusters was counted.

### Autoantibody detection

Autoantibodies were detected in serum samples using a NovaLite rat liver, kidney, and stomach multicomposite kit: “Composite tissue slides” (Innova Diagnostics). Briefly, blood was drawn from a facial blood vessel into a microtube precoated with 0.2 μl 0.5 M EDTA and spun down (4°C, 2000 x g, 15 min) to obtain the sera. Composite tissue slides were incubated with 1/40 sera at room temperature according to an established protocol (*60*) followed by detection with goat anti–mouse IgG (H+L) Alexa 555 (ThermoFischer Scientific). Images were acquired using a DM6000 microscope (Leica Microsystems). Quantification of autoantibodies was performed by measuring mean fluorescence intensity (MFI) at selected regions of interest corresponding to specific tissue areas using the ImageJ program (NIH). Data across the experiments was normalized according to the MFI background of a secondary antibody staining.

### Analysis of autoimmunity in the intestine

For the counting of Paneth cells (PC), mouse terminal ileums were isolated, cleared of feces, and fixed in 4% paraformaldehyde overnight. They were then placed in 70% ethanol overnight, dehydrated, and embedded in paraffin using the Leica HistoCore Pegasus Tissue Processor, and subsequently the Leica HistoCore Arcadia. Embedded ileums were longitudinally cut into 10 μm sections. Prior to staining, sections were blocked with 10% bovine serum albumin in PBS. To visualize PC, paraffin sections were stained with polyclonal anti-lysozyme antibody (Agilent/Dako; host: rabbit). Then, goat anti-rabbit IgG (H+L) Alexa 555-conjugated secondary antibody (ThermoFisher Scientific) was applied followed by DAPI staining. The average number of PCs per crypt was determined as previously described (*90*). For analysis of the colon weight/length ratio, mouse colons were isolated along with the cecum. Isolated tissue was stretched and the length from the cecum to rectum was measured. The colons were then separated from the cecums, cleaned of feces, and weighed. The ratio was calculated by dividing the weight of the colon (mg) by the length of the colon (mm).

### Statistical analysis

Statistical analysis and graphs were generated using Prism 5.0.4 and 10.4.0 software (GraphPad), except for scRNAseq, which was analyzed using the Seurat package, R v 4.0.2 (R core team 2020). When comparing two experimental groups, unpaired or paired Student’s t-test was used. One-way ANOVA with Tukey’s multiple comparison test was used to compare three or more experimental groups. When pairing between samples was applicable, repeated measures (RM) one-way ANOVA with Tukey’s multiple comparisons test was used. Survival curve statistics was analyzed using the log-rank (Mantel-Cox) test. In the scatter plots and BrdU lineage tracing plot, the mean ± SEM is shown. Median and quartiles are shown for some of the violin plots. Sample sizes, experimental replicates, and additional information such as type of normalization are provided in the figure legends. If p ≤ 0.05, it is considered statistically significant. In certain cases where we refer to a possible statistically significant trend, the abbreviation “ns” has been replaced by the exact p-value in the figures. The pipeline of scRNAseq analysis is described in the sections “Single-cell RNA sequencing” and “Bioinformatic analysis”.

## Supplementary Materials

Supplementary Figures S1-S8.

Table S1. List of antibodies

Data file S1, related to Figure S1D. Marker genes of thymic myeloid cell subsets.

Data file S2, related to Figure S1E. Marker genes of aDC subsets.

Data file S3, related to Figure 1D. DEGs between TdTOMATO^+^ and TdTOMATO^−^ DC1.

Data file S4. Source data.

Data file S5. Seurat object related to scRNAseq analysis from Fig. 1B

Data file S6. Seurat object related to scRNAseq analysis from Fig. S1B

## Acknowledgments

We would like to thank Z. Cimburek, M. Šíma, and Š. Kocourková from the Flow cytometry as well as the Genomics and Bioinformatics facilities of the Institute of Molecular Genetics of the Czech Academy of Sciences for their technical support. We would also like to thank R. S. Blumberg (Division of Gastroenterology, Department of Medicine, Brigham and Women’s Hospital, Harvard Medical School, Boston, MA) for providing *Defa6^iCre^* mice, V. Kořínek (IMG, Prague, Czech Republic) for *R26^TdTOMATO^* mice, and L. Klein (Ludwig Maximilian University, Munich, Germany) for *Adig^GFP^* and *Aire^−/–^*mice. We are grateful to E. Cosway, K. James, A. White, and S. Parnell (Institute of Immunology and Immunotherapy, University of Birmingham, Birmingham, UK) for advice regarding the use of composite tissue slides, BrdU lineage tracing experiment, and animal handling related to this experiment. We would like to thank V. Uleri and A. Neuwirth (IMG, Prague, Czech Republic) for their methodological advice. Finally, we would like to acknowledge K. Jelínková (Faculty of Science, Charles University, Prague, Czech Republic) for the illustration of mouse figures.

## Author contributions

JB, MV, and DF initiated the project and co-designed experiments. JB conducted the majority of experiments and wrote the manuscript. TB designed some experiments, performed bioinformatic analysis of scRNAseq and along with MV and JM edited the manuscript. DM, VS, KJ, EV, and KK conducted some experiments. OB co-designed and conducted autoantibody detection. JP co-designed and performed light sheet fluorescence microscopy experiment. VT, AČ, and MD provided technical support. JD provided guidance and methodological advice. JK and MK performed scRNAseq. PK and RS generated *R26^TdT-OVA^*mouse model. OŠ and JČ provided methodological advice and resources. ST and BM provided mouse models. GA co-designed BrdU lineage tracing experiment, provided methodological advice and resources. DF acquired funding, supervised research, and finalized the manuscript.

## Competing interests

The authors declare no competing interests.

### Data and materials availability

The scRNAseq data are available in the BioStudies database under the accession number E-MTAB-14319 (https://www.ebi.ac.uk/biostudies/arrayexpress/studies/E-MTAB-14319). Source data files for all figures and supplementary figures can be found in Data file S4. All data needed to evaluate the conclusions of this study are present in the paper or in Supplementary Materials.

## Supplementary figures

**Figure S1.**
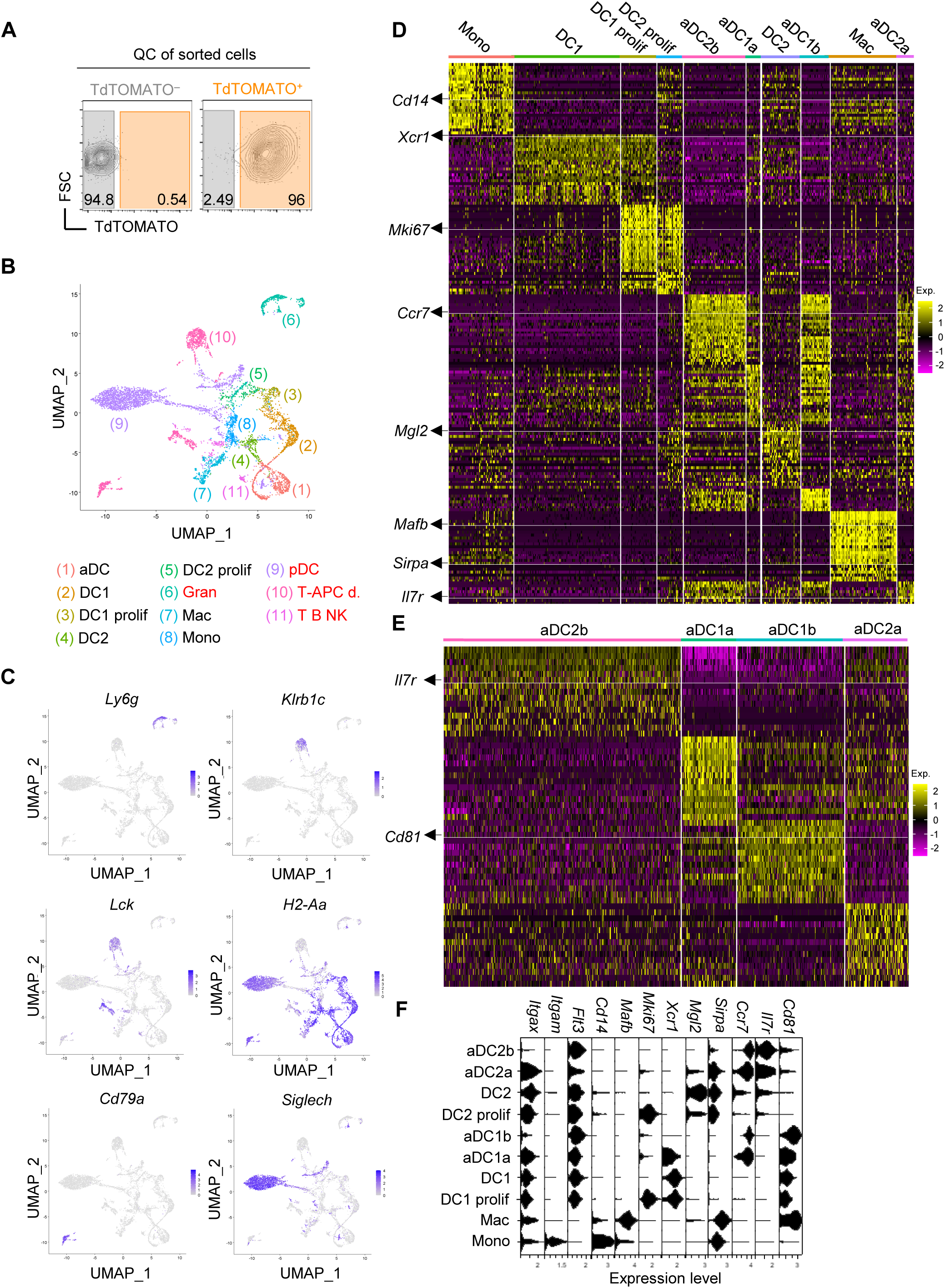
Single-cell RNA sequencing of thymic myeloid APCs. **(A)** The quality control contour plots showing the expression level of TdTOMATO in sorted TdTOMATO^−^ and TdTOMATO^+^ thymic myeloid cells to perform scRNAseq. **(B)** UMAP of scRNAseq of all sorted and annotated thymic myeloid cells. The red font marks subsets that were excluded from the analysis. DC=conventional DC, aDC=activated DC, Gran=Granulocytes, Mac=Macrophages, Mono=Monocytes, pDC=plasmacytoid DC, T-APC d.=doublets of T cells and antigen-presenting cells, T B NK=T, B, and NK cells, prolif=proliferating. **(C)** UMAP featureplots showing the expression of marker genes of cell subsets that were excluded from scRNAseq analysis. **(D)** Heatmap showing up to 25 of the top marker genes of each subset from scRNAseq annotated in Fig. 1B. Heatmap color scale depicts average log2 fold change. For better discernibility of the expression profile across all subsets annotated in Fig 1B, the selected marker genes are highlighted by arrows and underlined with a white line. **(E)** Heatmap showing up to 15 of the top marker genes of each aDC subset from scRNAseq. Heatmap color scale depicts average log2 fold change. **(F)** Violin plots show the expression of selected marker genes of subsets annotated in Fig. 1B.

**Figure S2.**
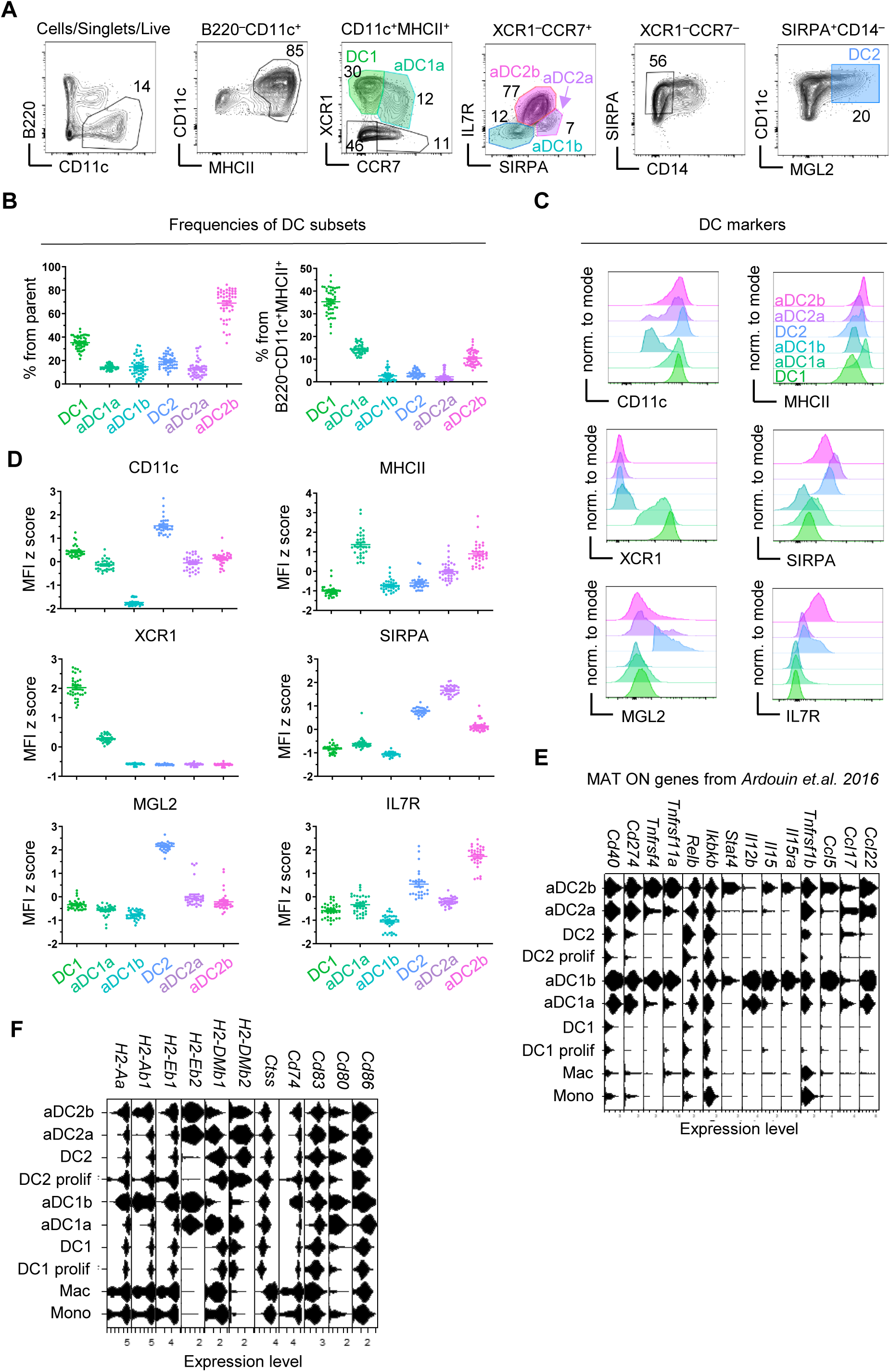
The expression of marker and antigen presentation-associated genes by thymic DCs. **(A)** Flow cytometry gating strategy of thymic DCs based on scRNAseq of thymic myeloid cells. DCs were gated as B220^−^CD11c^+^MHCII^+^ and further distinguished into XCR1^+^CCR7^−^ DC1 and XCR1^+^CCR7^+^ aDC1a. XCR1^−^CCR7^+^ DCs were further separated into SIRPA^High^IL7R^Low^ aDC2a, SIRPA^Low^IL7R^High^ aDC2b and SIRPA^−^IL7R^−^ aDC1b. XCR1^−^CCR7^−^ cells were then gated as SIRPA^+^MGL2^+^CD14^−^ DC2. Color code of thymic DC subsets defined here is used across all figures. **(B)** Frequencies of DC subsets within parent populations (left panel) and within all B220^−^ CD11c^+^MHCII^+^ cells (right panel) (mean ± SEM, *n*=45-49 mice from 7 independent experiments) gated as in Fig. S2A. **(C)** Histograms depicting the protein expression of DC markers in DC subsets as defined in Fig. S2A. **(D)** MFI z scores of DC markers within DC subsets related to Fig. S2C (mean ± SEM, *n*=29-33 mice from 5 independent experiments). **(E)** Violin plots show the expression of MAT ON genes taken from *Ardouin et.al. 2016* within cell subsets annotated in Fig. 1B. **(F)** Violin plots show the expression of genes involved in MHCII presentation within cell subsets annotated in Fig. 1B.

**Figure S3.**
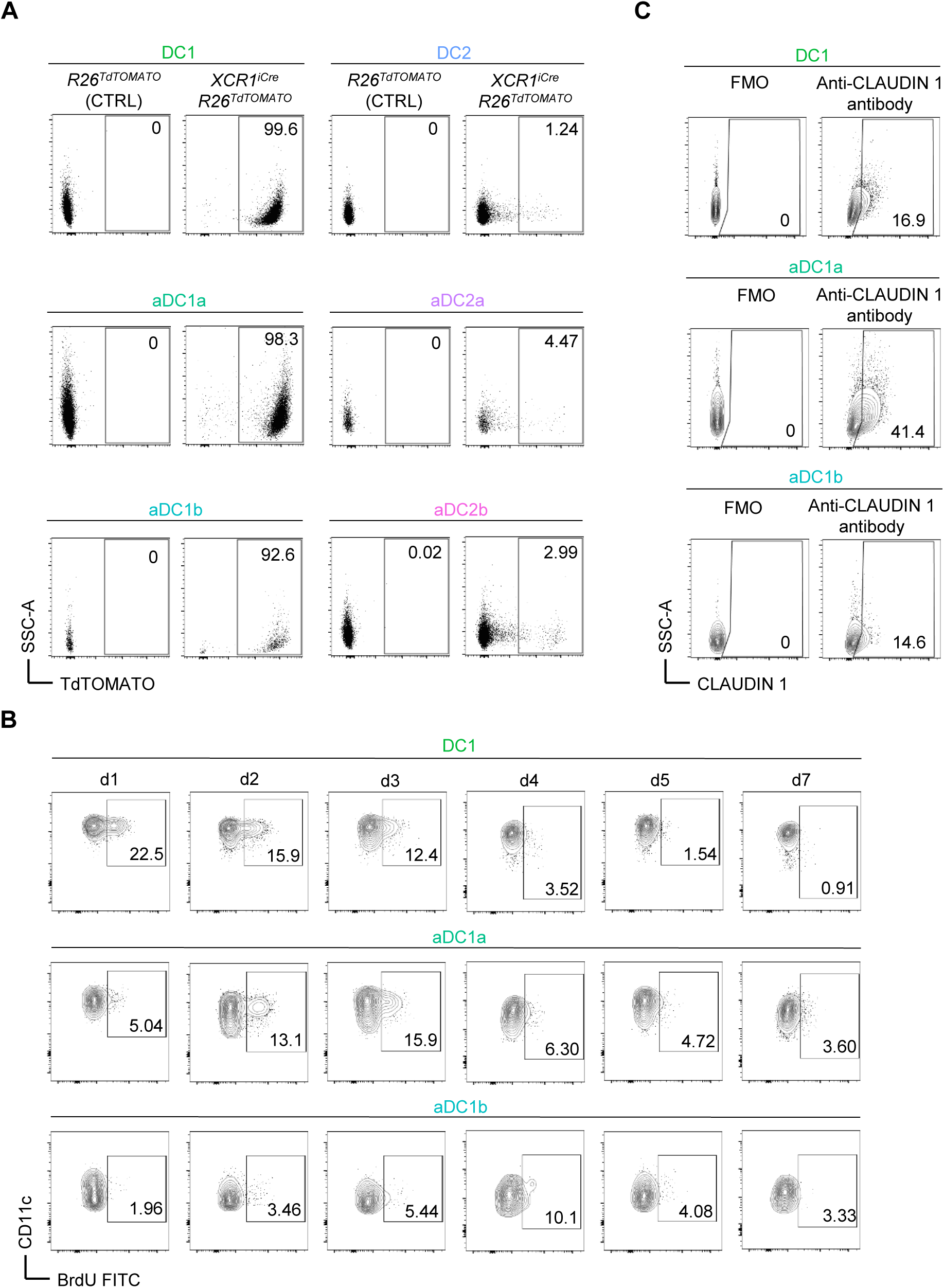
Thymic DC1 lineage maturation yields three successive stages. **(A)** Representative flow cytometry plots show TdTOMATO expression within DC1 subsets from mouse model in Fig. 2A. **(B)** Representative flow cytometry plots show the frequency of BrdU^+^ cells within DC1 subsets on indicated days after the BrdU administration related to Fig. 2D. **(C)** Representative flow cytometry plots show CLAUDIN 1 positivity within DC1 subsets related to Fig. 2E. FMO controls are shown.

**Figure S4.**
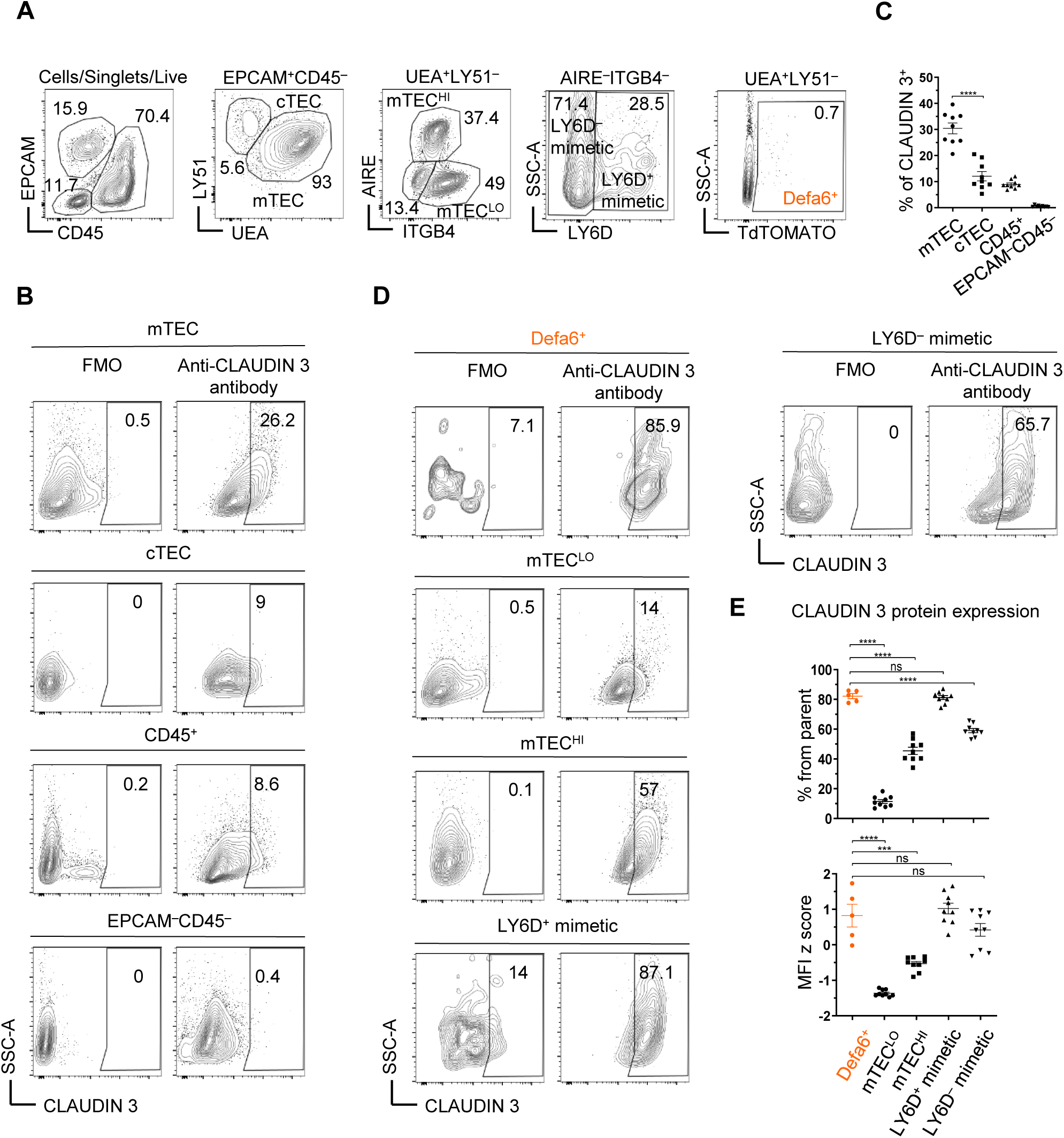
Defa6^+^ mTECs and keratinocyte mimetic cells express a ligand of Claudin 1. **(A)** Flow cytometry gating strategy of mTEC subsets. Thymic epithelial cells were gated as EPCAM^+^CD45^−^ and further distinguished into LY51^+^UEA^−^ cortical thymic epithelial cells (cTEC) and LY51^−^UEA^+^ mTECs. The latter were further separated into AIRE^+^ITGB4^−^ mTEC^HI^, AIRE^−^ ITGB4^+^ mTEC^LO^ and double negative cells which were comprised of keratinocyte mimetics (LY6D^+^) and other mimetic cells (LY6D^−^). Defa6^+^ mTECs are color coded in orange denoting TdTOMATO positivity. **(B)** Representative flow cytometry plots show CLAUDIN 3 positivity within mTECs, cTECs, CD45^+^ cells (hematopoietic cells) and EPCAM^−^CD45^−^ cells (fibroblasts, endothelial cells etc.). FMO controls are shown. **(C)** The frequency of CLAUDIN 3^+^ cells within cell populations from Fig. S4B (mean ± SEM, *n*=9 mice from 3 independent experiments). **(D)** Representative flow cytometry plots show CLAUDIN 3 positivity within individual mTEC subsets gated as in Fig. S4A. FMO controls are shown. **(E)** The frequency of CLAUDIN 3^+^ cells and MFI z score of CLAUDIN 3 expression within mTEC subsets from Fig. S4D (mean ± SEM, *n*=5-9 mice from 3 independent experiments). Statistical analysis in C and E was performed using unpaired, two-tailed Student’s t-test, p ≤ 0.001***, p < 0.0001 = ****, ns = not significant.

**Figure S5.**
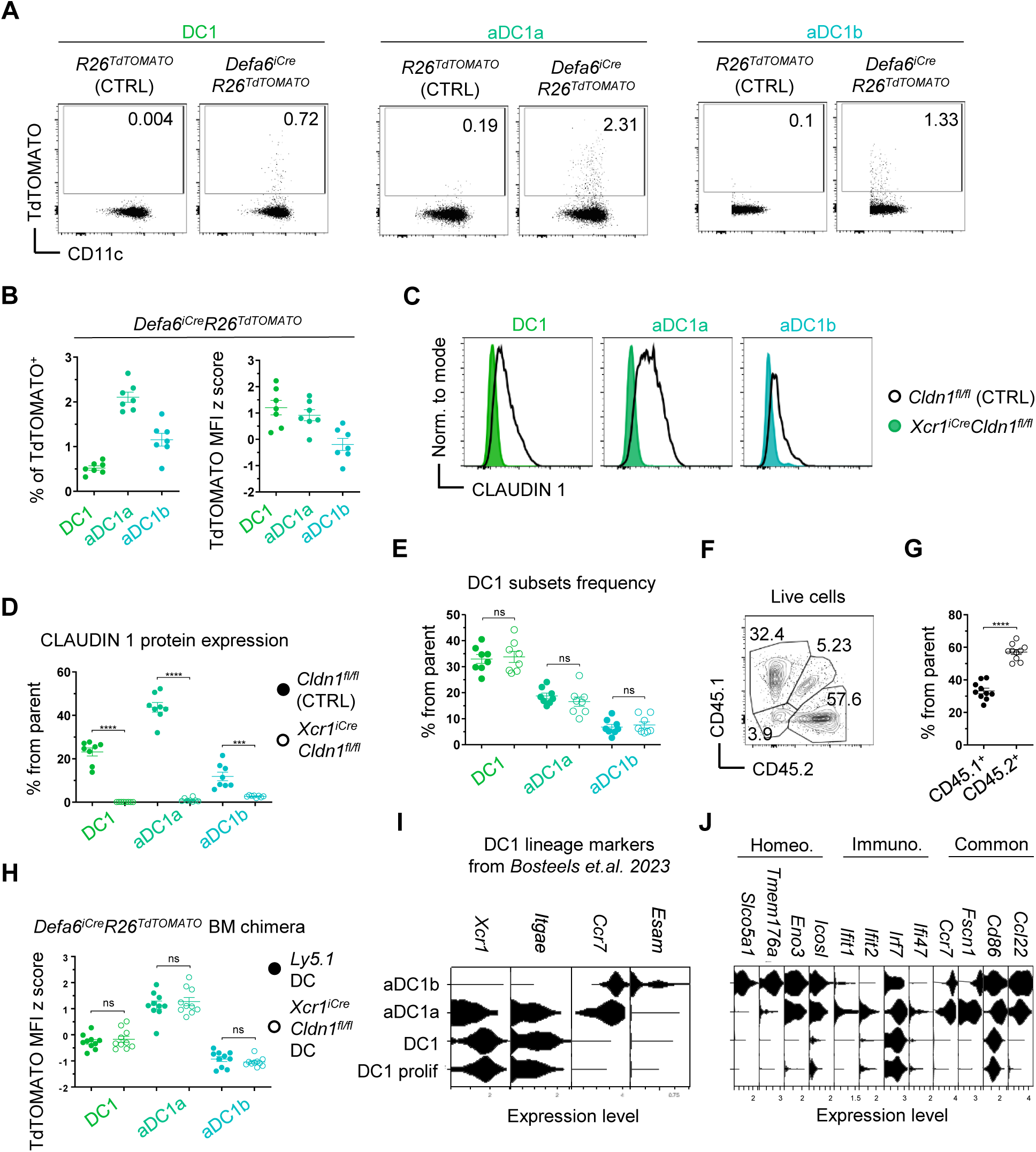
Mouse models to study the role of Claudin in CAT. **(A)** Representative flow cytometry plots show the acquisition of TdTOMATO by thymic DC subsets from *Defa6^iCre^R26^TdTOMATO^* mice. FMO control is shown (*R26^TdTOMATO^*). **(B)** Frequency of TdTOMATO^+^ cells and MFI z score of TdTOMATO expression within DC subsets from Fig. S5A (mean ± SEM, *n*=7 mice from 2 independent experiments). **(C)** Histograms show CLAUDIN 1 expression within thymic DC1 subsets from *Cldn1^fl/fl^* (control; black) and *XCR1^iCre^Cldn1^fl/fl^* (varying shades of green) mice. **(D)** Frequency of CLAUDIN 1^+^ cells within DC1 subsets related to Fig. S5C (mean ± SEM, *n*=8 mice from 3 independent experiments). **(E)** Frequency of DC1 lineage subsets within parent populations from *Cldn1^fl/fl^* (control; solid circle) and *XCR1^iCre^Cldn1^fl/fl^* (empty circle) mice (mean ± SEM, *n*=8 mice from 3 independent experiments). **(F)** Representative flow cytometry plot shows the reconstitution of competitive BM chimeras from Fig. 3A. **(G)** Quantification of BM reconstitution from Fig. S5E (mean ± SEM, *n*=17 mice from 4 independent experiments). **(H)** MFI z score of TdTOMATO expression within Claudin 1-sufficient and -deficient DC1 subsets from Fig. 3B-C (mean ± SEM, *n*=10 mice from 3 independent experiments). **(I)** Violin plots from scRNAseq analysis (Fig. 1) show the expression of DC1 lineage markers used for flow cytometry gating strategy of DC1 subsets in *Bosteels et.al. 2023*. **(J)** Violin plots from scRNAseq analysis (Fig. 1) show the expression of selected marker genes of homeostatic (Homeo.) and immunogenic (Immuno.) maturation and genes associated with both maturation programs (Common) from *Bosteels et.al. 2023*. The subsets analyzed are the same as in Fig. S5I. Statistical analysis in G and H was performed using paired while D and E was by unpaired, two-tailed Student’s t-test, p ≤ 0.001***, p < 0.0001 = ****, ns = not significant.

**Figure S6.**
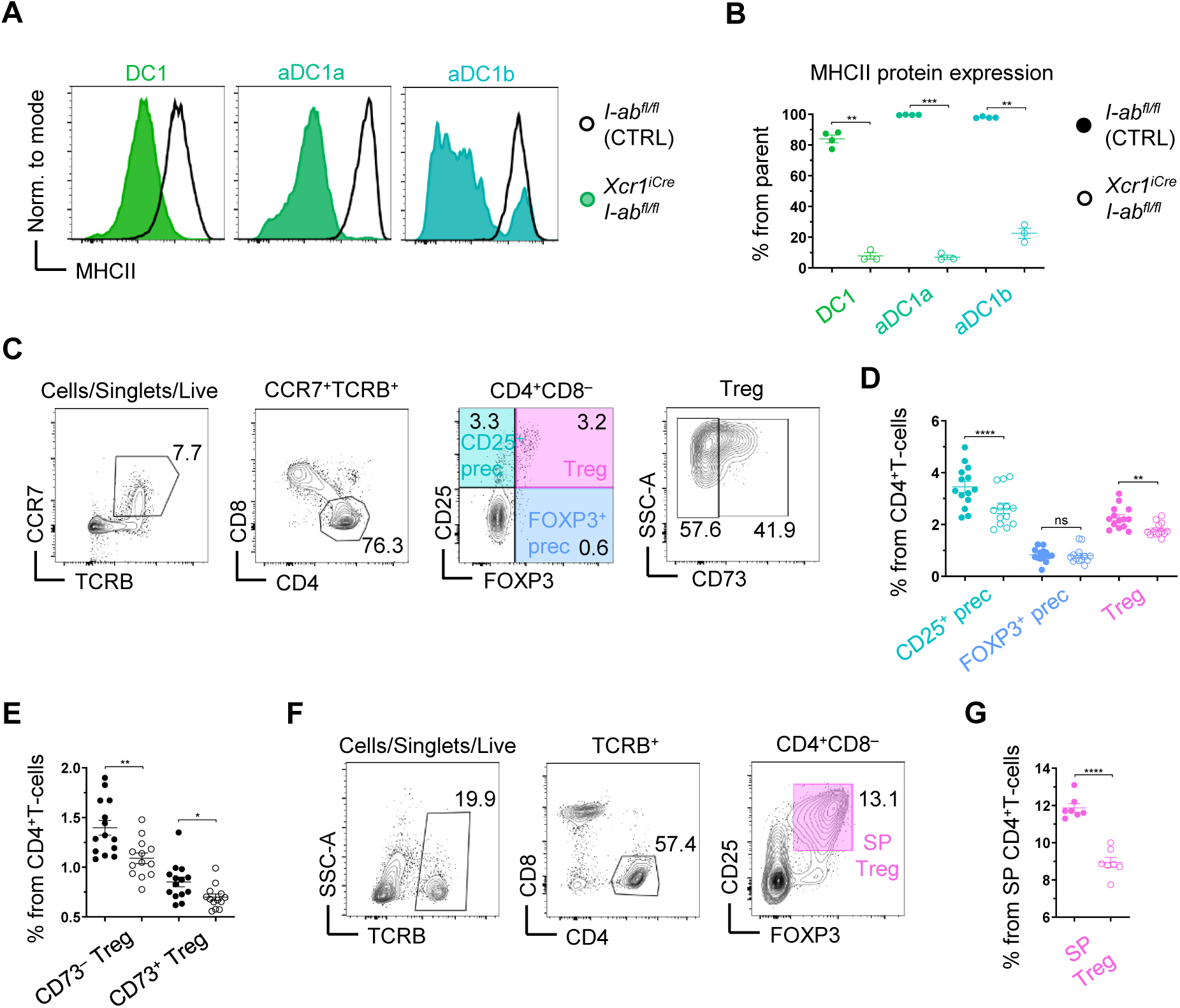
Antigen presentation by thymic DC1 lineage is required for Treg selection. **(A)** Histograms showing MHCII expression within DC1 lineage subsets from *I-ab^fl/fl^* (control; black) and *XCR1^iCre^I-ab^fl/fl^* (varying shades of green) mice. **(B)** Frequency of MHCII^+^ cells within DC1 subsets from *I-ab^fl/fl^* (control; solid circle) and *XCR1^iCre^I-ab^fl/fl^* (empty circle) mice, related to Fig. S6A (mean ± SEM, *n*=3-4 mice from 2 independent experiments). **(C)** Flow cytometry gating strategy of thymic T cells. T cells were gated as CCR7^+^TCRB^+^ to analyze medullary T cells only. These were gated as CD4^+^ (helper T cells) which were further separated into CD25^+^FOXP3^−^ Treg precursors (CD25^+^ prec), CD25^−^FOXP3^+^ Treg precursors (FOXP3^+^ prec) and CD25^+^FOXP3^+^ Treg. Recirculating Tregs were distinguished from the newly generated as CD73^+^. Color code of thymic T cell subsets is defined here. **(D)** Frequency of Tregs and their precursors within CD4^+^ T cells isolated from thymi of *I-ab^fl/fl^* (control; solid circle) and *XCR1^iCre^I-ab^fl/fl^* (empty circle) mice (mean ± SEM, *n*=13-14 mice from 4 independent experiments). **(E)** Frequency of CD73^−^ and CD73^+^ cells within Treg subset from thymi of *I-ab^fl/fl^* (control; solid circle) and *XCR1^iCre^I-ab^fl/fl^* (empty circle) mice (mean ± SEM, *n*=13-14 mice from 4 independent experiments). **(F)** Flow cytometry gating strategy of splenic T cells which were gated as TCRB^+^CD4^+^ and further separated into Tregs (SP Treg) based on the expression of CD25 and FOXP3. Color code of splenic Tregs is shown. **(G)** Frequency of SP Tregs within CD4^+^ T cells from *I-ab^fl/fl^* (control; solid circle) and *XCR1^iCre^I-ab^fl/fl^* (empty circle) mice, related to S6G (mean ± SEM, *n*=7 mice from 2 independent experiments). Statistical analysis in B, D, E, and G was performed using unpaired, two-tailed Student’s t-test, p ≤ 0.05 = *, p ≤ 0.01 = **, p ≤ 0.001***, p < 0.0001 = ****, ns = not significant.

**Figure S7.**
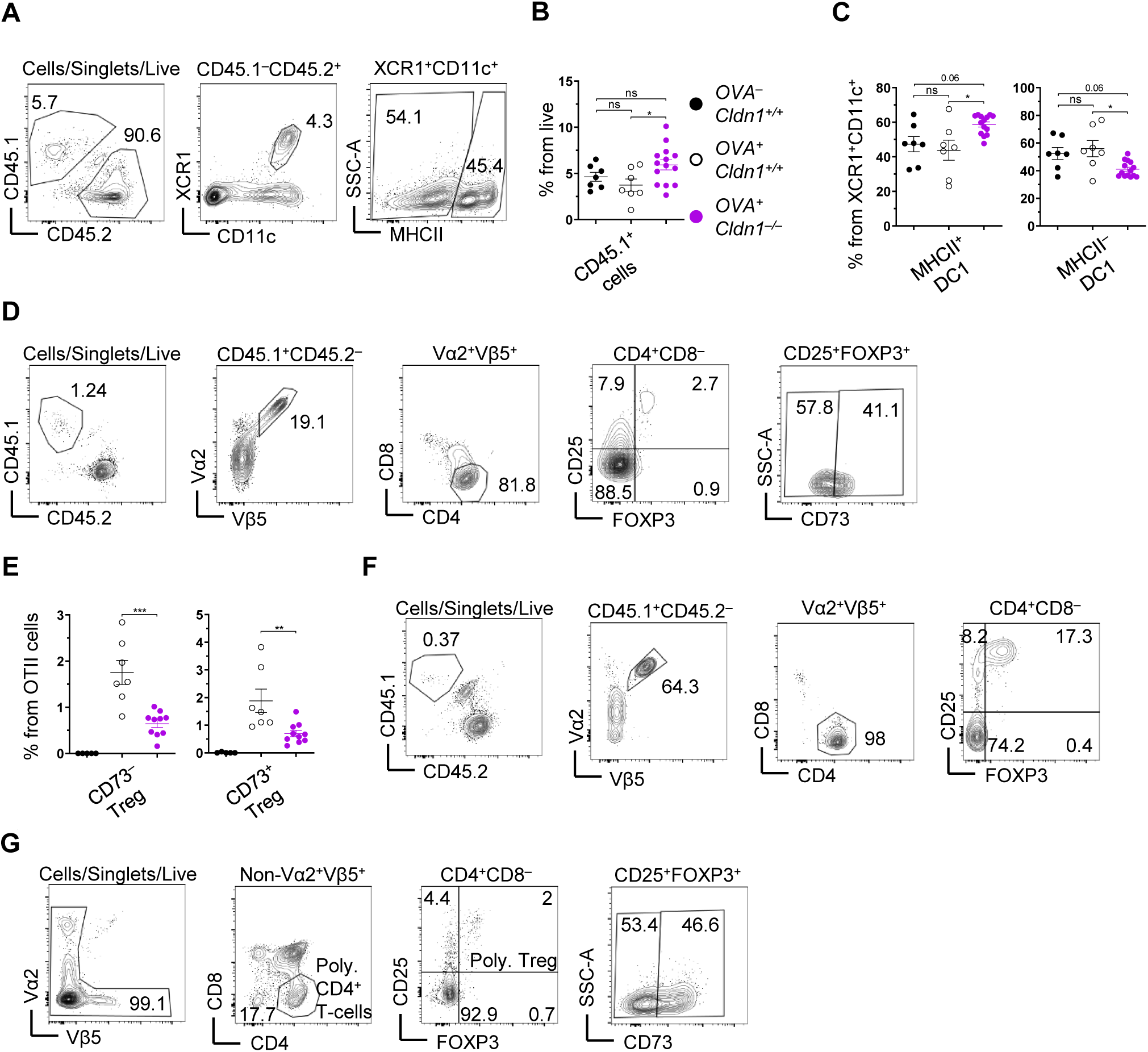
*Defa6^iCre^R26^TdT-OVA^* mouse as a tool to study relevance of Claudin 1 in the selection of TRA-specific T cells. **(A)** Flow cytometry gating strategy to analyze reconstitution of competitive BM chimeras from Fig. 5B-C. Note that Cldn1^+/+^MHCII^−/–^, Cldn1^−/–^MHCII^+/+^ and Cldn1^+/+^MHCII^+/+^ BM were all CD45.2^+^; thus, we quantified the reconstitution of each particular mixture of these BM based on the expression of MHCII within DC1 lineage. **(B)** Quantification of reconstitution of CD45.1^+^ BM (mean ± SEM, *n*=7-14 mice from 3 independent experiments). Color code is the same as described in Fig. 5C. **(C)** Quantification of reconstitution of BM comprised of MHCII-sufficient (left panel) and MHCII-deficient (right panel) thymic DC1 lineage (mean ± SEM, *n*=7-14 mice from 3 independent experiments). Color code as in Fig. S7B. **(D)** Flow cytometry gating strategy of OTII thymic T cells. CD45.1^+^ cells were gated as Vα2^+^Vβ5^+^ to obtain OTII cells. These cells were then gated as CD4^+^ and separated into the same populations as in Fig. S6D and conventional CD25^−^FOXP3^−^OTII cells. **(E)** Frequency of newly generated CD73^−^ (left panel) and recirculating CD73^+^ (right panel) cells within OTII Tregs from thymi of competitive BM chimeras described in Fig. 5B-C (mean ± SEM, *n*=5-10 mice from 2 independent experiments). Color code as in Fig. S7B. **(F)** Flow cytometry gating strategy of OTII cells from skin draining lymph nodes. CD45.1^+^ OTII cells were gated as Vα2^+^Vβ5^+^. These were then gated as CD4^+^ and separated into CD25^+^FOXP3^+^ OTII Tregs and conventional CD25^−^FOXP3^−^ OTII cells. **(G)** Flow cytometry gating strategy of developing polyclonal T-cells from thymus. CD4^+^ polyclonal T-cells were pregated as non-Vα2^+^Vβ5^+^ to leave out OTII cells from further analysis. These cells were further gated as polyclonal Tregs which were further distinguished into newly generated (CD73^−^) and recirculating (CD73^+^). Statistical analysis in B and C was performed using one-way ANOVA with Tukeýs multiple comparisons test and in E was performed using unpaired, two-tailed Student’s t-test, p ≤ 0.05 = *, p ≤ 0.01 = **, p ≤ 0.001***, ns = not significant.

**Figure S8.**
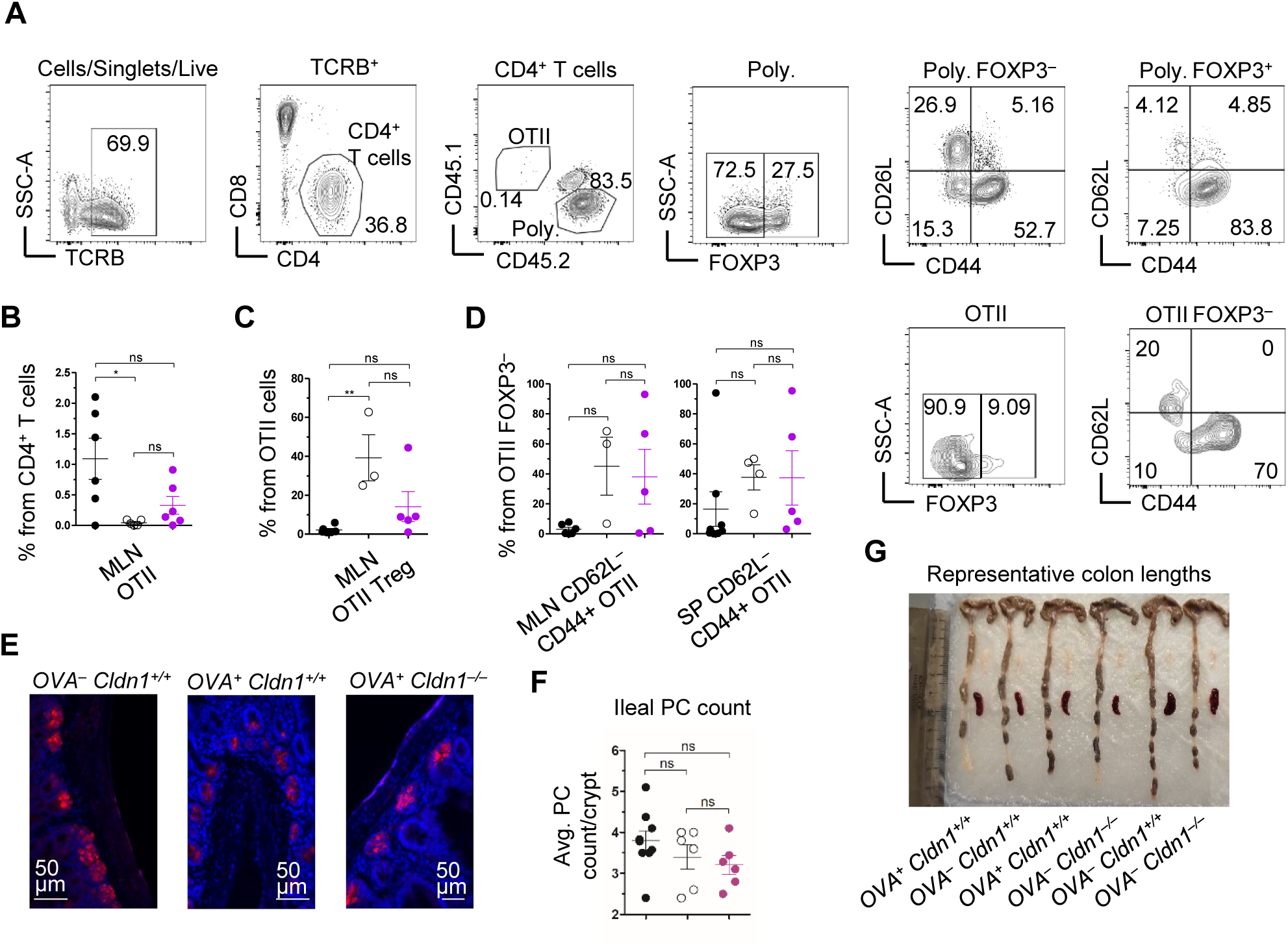
Autoimmune manifestations of Claudin 1-deficient bone marrow chimeras. **(A)** Representative flow cytometry gating strategy of activated T cells from mesenteric lymph node (MLN) and spleen. CD4^+^ T cells were gated as TCRB^+^CD4^+^CD8^−^ and further distinguished using congenic markers to OTII (CD45.1^+^CD45.2^−^) and polyclonal (Poly.; CD45.1^−^CD45.2^+^) cells. Both OTII and polyclonal cells were further distinguished into conventional T cells (FOXP3^−^) and Tregs (FOXP3^+^). These populations were analyzed for CD62L and CD44 expression, with CD62L^−^CD44^+^ cells representing effector memory T cells. **(B-C)** Frequency of OTII cells within CD4^+^ T cells (B) and OTII Tregs within OTII cells (C) isolated from MLN of BM chimeras from Fig. 5B-C around 40 weeks post BM transfer, gated asi in Fig. S8A (mean ± SEM, *n*=3-6 mice from 2 independent experiments). **(D)** Frequency of effector memory CD62L^−^CD44^+^ cells within conventional OTII cells (FOXP3^−^) isolated from MLN (left panel) and spleen (right panel) of BM chimeras from Fig. 5B-C around 40 weeks post BM transfer, gated as in Fig. S8A (mean ± SEM, *n*=3-8 mice from 2 independent experiments). **(E)** Representative immunofluoresence images of ileal Paneth cells (PC) stained by lysozyme (red) and DAPI (blue) of BM chimeras from Fig. 5B-C around 40 weeks post BM transfer. **(F)** Quantification of average PC count per crypt within ileums of BM chimeras from Fig. 5B-C around 40 weeks post BM transfer related to Fig. S8E (mean ± SEM, *n*=6-9 mice from 2 independent experiments). **(G)** Representative colon lengths of BM chimeras from Fig. 5B-C around 40 weeks post BM transfer. Statistical analysis in B, C, D and F was performed using one-way ANOVA with Tukeýs multiple comparisons test, p ≤ 0.05 = *, p ≤ 0.01 = **, ns = not significant.

