## Supplemental Table S1 for "Claudin 1-mediated positioning of DC1 to mTECs is essential for antigen transfer-coupled DC1 maturation and maintenance of central tolerance"

| Antibody | Manufacturer | Clone | Catalogue no. | Dilution |
| --- | --- | --- | --- | --- |
| Anti-BrdU-FITC | BD | BrdU Flow Kit | cat# 559619 | 1:100 |
| Anti-mouse AIRE-AF488 | eBioscience | 5H12 | cat# 53-5934-82 | 1:200 |
| Anti-mouse CCR7-BV605 | BioLegend | 4B12 | cat# 120125 | 1:100 |
| Anti-mouse CD11c-APC/Cy7 | BioLegend | N418 | cat# 117324 | 1:200 |
| Anti-mouse CD11c-Biotin | eBioscience | N418 | cat# 13-0114-82 | 1:100 |
| Anti-mouse CD14-FITC | BioLegend | Sa2-8 | cat# 123308 | 1:100 |
| Anti-mouse CD25-PE/Cy7 | BioLegend | PC61 | cat# 102016 | 1:200 |
| Anti-mouse CD301b (MGL2)-PE/Cy7 | BioLegend | URA-1 | cat# 146807 | 1:200 |
| Anti-mouse CD326 (EPCAM)-BV785 | BioLegend | G8.8 | cat# 118245 | 1:1000 |
| Anti-mouse CD44-A700 | BioLegend | IM7 | cat# 103026 | 1:200 |
| Anti-mouse CD45.1-BV605 | BioLegend | 104 | cat# 110738 | 1:100 |
| Anti-mouse CD45.1-PerCP/Cy5.5 | BioLegend | A20 | cat# 110727 | 1:100 |
| Anti-mouse CD45.2-BUV395 | BD | 104 | cat# 564616 | 1:100 |
| Anti-mouse CD45-BV605 | BioLegend | 30-F11 | cat# 103155 | 1:100 |
| Anti-mouse CD45-FITC | BioLegend | 30-F11 | cat# 103108 | 1:100 |
| Anti-mouse CD4-BV785 | BioLegend | GK1.5 | cat# 100453 | 1:300 |
| Anti-mouse CD73-PB | Biolegend | TY/11.8 | cat# 127211 | 1:200 |
| Anti-mouse CD8a-Biotin | BioLegend | 53-6.7 | cat# 100704 | 1:200 |
| Anti-mouse FOXP3-APC | eBioscience | FJK-16s | cat# 17-5773-82 | 1:200 |
| Anti-mouse FOXP3-Percp Cy5.5 | eBioscience | FJK-16s | cat# 45-5773-82 | 1:200 |
| Anti-mouse I-A/I-E-BUV737 | BD | M5/114.15.2 | cat# 748845 | 1:800 |
| Anti-mouse I-A/I-E-BV711 | BioLegend | M5/114.15.2 | cat# 107625 | 1:700 |
| Anti-mouse IL7R-APC | Biolegend | A7R34 | cat# 135012 | 1:100 |
| Anti-mouse ITGB4-Percp Cy5.5 | BioLegend | 364-11A | cat# 123614 | 1:100 |
| Anti-mouse Ly-51-PE/Cy7 | Biolegend | 6C3 | cat# 108314 | 1:200 |
| Anti-mouse Ly6D-AF450 | eBioscience | 49-H4 | cat# 48-5974-80 | 1:200 |
| Anti-mouse SIRPA-BUV563 | BD | P84 | cat# 741349 | 1:150 |
| Anti-mouse SIRPA-Percp eFluor 710 | eBioscience | P84 | cat# 46-1721-82 | 1:150 |
| Anti-mouse TCRB-BV421 | Biolegend | H57-597 | cat# 109230 | 1:150 |
| Anti-mouse TCRB-FITC | Biolegend | H57-597 | cat# 109206 | 1:150 |
| Anti-mouse Vα2-PE | BioLegend | B20.1 | cat# 127808 | 1:150 |
| Anti-mouse Vβ5.1/Vβ5.2-FITC | BD | MR9-4 | cat# 51-01354L | 1:75 |
| Anti-mouse/human CD11b-Biotin | Biolegend | M1/70 | cat# 101204 | 1:100 |
| Anti-mouse/human B220-BV785 | BioLegend | RA3-6B2 | cat# 103246 | 1:100 |
| Anti-mouse/human/dog CLAUDIN 1 | eBioscience | Polyclonal | cat# 51-9000 | 1:100 |
| Anti-mouse/human/dog CLAUDIN 3 | eBioscience | Polyclonal | cat# 34-1700 | 1:100 |
| Anti-mouse/rat XCR1-BV421 | BioLegend | ZET | cat# 148216 | 1:200 |
| FC Block TruStain FcX™ PLUS | BioLegend | S17011E | cat# 156604 | 1:100 |
| Fixable Viability Dye-eFluor 506 | eBioscience | - | cat# 65-0866-14 | 1:300 |
| Goat anti-mouse-AF555 | eBioscience | - | cat# A-21422 | 1:500 |
| Goat anti-rabbit-AF488 | eBioscience | - | cat# A-11034 | 1:500 |
| Goat anti-rabbit-AF555 | eBioscience | - | cat# A-21429 | 1:500 |
| Goat anti-rabbit-AF647 | eBioscience | - | cat# A-21244 | 1:500 |
| Rabbit anti-human lysozyme | Agilent/Dako | Polyclonal | cat# F0372 | 1:500 |
| Streptavidin-APC/Cy7 | BioLegend | - | cat# 405208 | 1:500 |
| UEA I-Biotin | Vector Labs | - | cat# B-1065-2 | 1:500 |

**Table S1. List of antibodies**
